# Morphogenesis and development of human telencephalic organoids in the absence and presence of exogenous ECM

**DOI:** 10.1101/2022.12.06.519271

**Authors:** Catarina Martins-Costa, Vincent Pham, Jaydeep Sidhaye, Maria Novatchkova, Angela Peer, Paul Möseneder, Nina S. Corsini, Jürgen A. Knoblich

## Abstract

Establishment and maintenance of apical-basal polarity is a fundamental step in brain development, instructing the organization of neural progenitor cells (NPCs) and the developing cerebral cortex. Particularly, basally located extracellular matrix (ECM) is crucial for this process. In vitro, epithelial polarization can be achieved via endogenous ECM production, or exogenous ECM supplementation. While neuroepithelial development is recapitulated in cerebral organoids, the effects of different ECM sources in tissue morphogenesis remain unexplored. Here, we show that exposure to exogenous ECM at early neuroepithelial stages causes rapid tissue polarization and complete rearrangement of neuroepithelial architecture within 3 days. In unexposed cultures, endogenous ECM production by NPCs results in gradual polarity acquisition over an extended time. After the onset of neurogenesis, tissue architecture and neuronal differentiation are largely independent of the initial ECM source. These results advance the knowledge on neuroepithelial biology in vitro, with a focus on mechanisms of exogenously- and endogenously-guided morphogenesis. They demonstrate the self-sustainability of neuroepithelial cultures by endogenous processes, prompting an urgent reassessment of indiscriminate use of exogenous ECM in these model systems.

## Introduction

Epithelial morphogenesis is an essential step in the development of several organs, including the brain (**Arai & Taverna**, 2017; **Hakanen et al**., 2019). In the central nervous system, crucial morphogenic steps take place during neurulation, when the neural plate gives rise to the neural tube (**Colas & Schoenwolf**, 2001; **Eom et al**., 2013; **Hakanen et al**., 2019). At that stage, polarized neuroepithelial cells present an apical domain adjacent to the fluid-filled ventricular lumen, and a basal domain at the outer neural tube surface (**Colas & Schoenwolf**, 2001). Analogous apical-basal organization is later found during corticogenesis, when the polarization of apical radial glia enables the stratified organization of other progenitor and neuronal populations across the cortical plate (**Arai & Taverna**, 2017). Impairment in these processes leads to severe neurodevelopmental defects (**Hakanen & Salminen**, 2015).

Polarization steps rely on coordinated signaling from neighboring cells and extracellular matrix (ECM) proteins. In particular, basement membrane proteins are cell surface-associated ECMs that line the basal surface of epithelial tissues, providing cues that initiate and maintain polarity (**Colognato et al**., 1999; **Datta et al**., 2011; **Henry & Campbell**, 1998; **Miner & Yurchenco**, 2004). In the brain, ECM production is carried out by different cell types, such as the meninges (**Decimo et al**., 2021) and neural progenitor cells (NPCs) themselves, and ECM signaling lies at the core of neurodevelopmental processes that differ between mice and humans (**Fietz et al**., 2012; **Florio & Huttner**, 2014; **Long et al**., 2018; **Namba et al**., 2019). Despite their central role in neurodevelopment (**Long & Huttner**, 2019), processes of ECM production and cell-ECM interactions remain understudied in cell culture models of the developing human brain.

Key features of neuroepithelium generation, morphogenesis, and differentiation can be recapitulated with brain organoids, three-dimensional models of developing human brain regions (**Eichmüller & Knoblich**, 2022; **Pasca**, 2018; **Quadrato & Arlotta**, 2017; **Sidhaye & Knoblich**, 2021). *In vitro* systems of various epithelial tissues are able to model aspects of endogenous morphogenesis with remarkable accuracy, but often lack endogenous ECM production, therefore requiring exogenous ECM supplementation (**Corsini & Knoblich**, 2022; **Inman & Bissell**, 2010; **Kratochvil et al**., 2019; **Simian & Bissell**, 2017). Accordingly, the vast majority of protocols used for brain organoid generation resorts to the early exposure of embryoid bodies (EBs) to exogenous ECM, which has been proposed as an adjuvant to the process of neuroepithelialisation (**Bhaduri et al**., 2020; **Eichmüller et al**., 2022; **Esk et al**., 2020; **He et al**., 2022; **Kelava et al**., 2022; **Paulsen et al**., 2022; **Qian et al**., 2018; **Velasco et al**., 2019; **Villa et al**., 2022). However, certain methodologies entirely forgo the addition of exogenous ECM (**Eiraku et al**., 2008; **Gordon et al**., 2021; **Sakaguchi et al**., 2015; **Yoon et al**., 2019). Currently, it is not known how these distinct experimental paradigms impact neuroepithelial development *in vitro*.

Here, we examined the effects of different modes of exogenous ECM exposure during human telencephalic organoid development. We resorted to Matrigel, an ECM preparation extracted from murine Engelbreth-Holm-Swarm sarcomas (**Orkin et al**., 1977) and mainly composed of ECM proteins - including laminin (60%), collagen IV (30%), entactin (8%), fibronectin, and heparan sulfate proteoglycan - and growth factors (**Corning Incorporated Life Sciences**, 2016). Matrigel was used in the formation of early 3D organoid-like cultures, such as mammary gland alveolar structures (**Barcellos-Hoff et al**., 1989; **Simian & Bissell**, 2017), and later to support *in vitro* culture of organoids from intestinal (**Sato et al**., 2009), brain (**Lancaster et al**., 2013), and other epithelial tissues (**Boretto et al**., 2017; **Dorrell et al**., 2014; **Eiraku et al**., 2011; **Gurumurthy et al**., 2022; **Huch et al**., 2013; **Jeong et al**., 2021; D. **Kim et al**., 2021; **Nakano et al**., 2012; **Nie et al**., 2017; **Stange et al**., 2013; **Turco et al**., 2017). Additionally, to assess the intrinsic ability of the neuroepithelium to endogenously produce ECM and undergo self-organization, we tested an experimental setup without any exogenous supplementation. The analysis was focused on early morphogenesis at pre-neurogenic stages, identity and organization of neural rosettes during the generation of deep-layer excitatory neurons, and long-term neuronal differentiation after the emergence of upper-layer excitatory neurons. We found that, while early stages of organoid development are remarkably different depending on the presence or absence of exogenous ECM, later stages are largely indistinguishable between the two conditions. With this systematic characterization, we have generated new insight into how the ECM influences neuroepithelial development *in vitro*, by exogenously- or endogenously-guided processes.

## Results

To evaluate how ECM supplementation influences early organoid development, we supplied exogenous ECM (exECM) in the form of Matrigel at the beginning of neuroepithelialisation (day 10, D10). Matrigel application was done either as a solid droplet that provides long-term exposure to a polymerized network of ECM proteins (droplet embedding, exECM^+D^) (**Lancaster et al**., 2017), or as transient dissolution in the culture medium (concentration of 2%V/V) from D10 to D13 (liquid embedding, exECM^+L^). These protocols were compared to one without exposure to exogenous ECM (exECM^-^) (**Materials and Methods**, **Fig. 1A**). To specifically investigate the effects on the dorsal telencephalon, which has a well-studied apical-basal polarity, we provided a three-day pulse of GSK3β-inhibitor CHIR99021 from D13 to D16, activating the WNT pathway and guiding neuroepithelial differentiation as previously described (**Lancaster et al**., 2017). Four human pluripotent stem cell lines (hPSCs) from healthy donors were used - one embryonic stem cell line (ESCs; H9), and three induced pluripotent stem cell lines (iPSCs; #1, #2 and #3; details in **Materials and Methods**). Analysis timepoints corresponded to important milestones in organoid development (**Fig. 1A**), starting with different stages of neuroepithelial morphogenesis, a process strongly influenced by ECM signaling.

**Figure 1.**
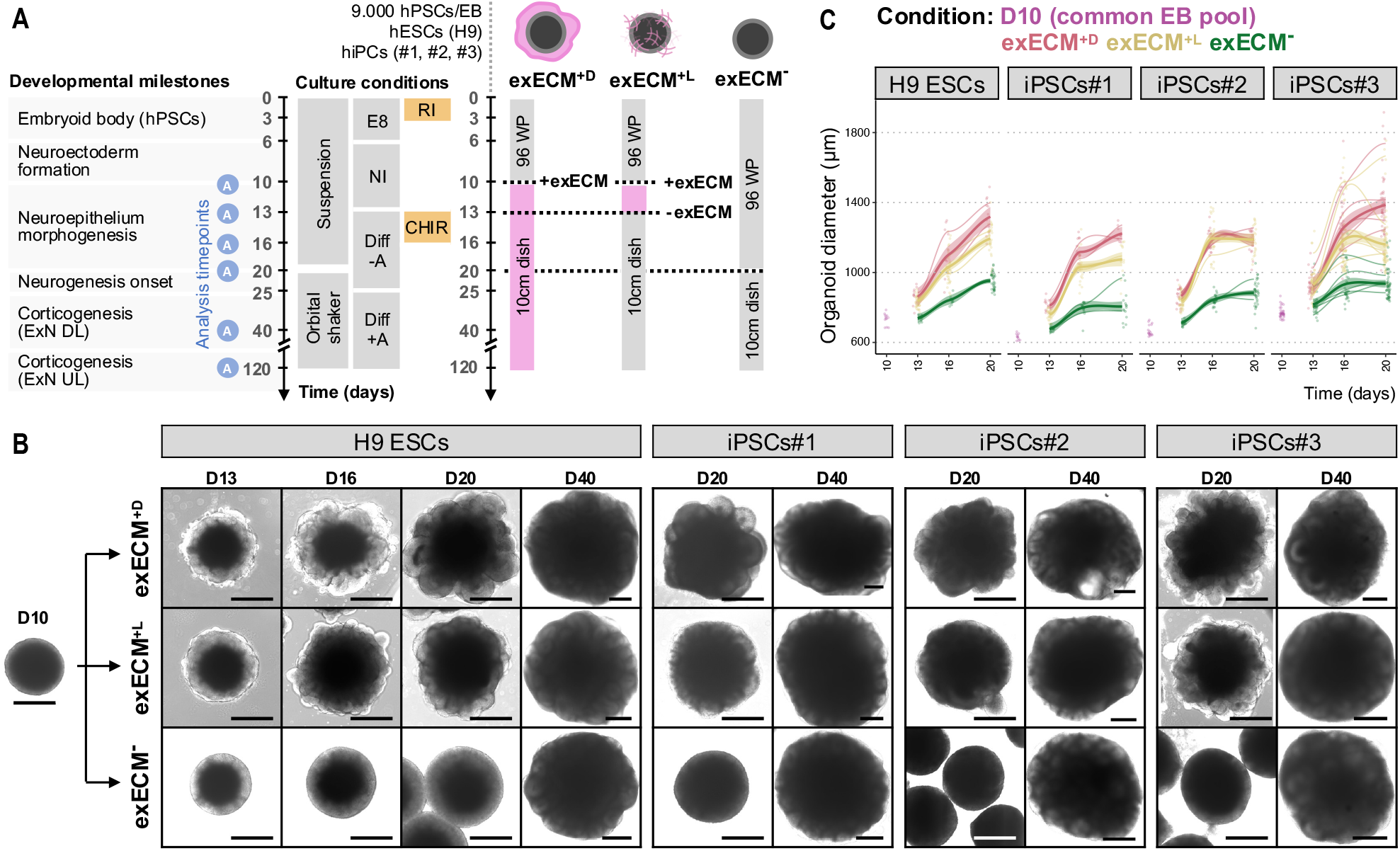
Exogenous ECM exposure influences organoid morphology and growth in the first 20 days of development. Summary of the three organoid protocols used (see Materials and Methods), timepoints of analysis, and relevant developmental milestones. A common pool of EBs was split into three experimental conditions at D10: embedding in a droplet of Matrigel (exECM^+D^), transient addition of Matrigel to the culture medium from D10 to D13 at a concentration of 2%V/V (exECM^+L^), and no exposure to any exogenous ECM (exECM^-^). Experiments were performed with 4 human pluripotent stem cell lines (hPSCs) and analysed at D10, D13, D16, D20, D40 and D120, when key developmental processes could be examined. (B) Organoid morphology in the first 40 days of culture. Note the appearance of tissue budding in exECM^+D^ and exECM^+L^ at D16, and the comparable organoid morphology across conditions at D40. Scale bars: 500 μm. (C) Organoid diameter in the first 20 days of culture. For each condition and cell line, datapoints were fit to a smoothed trend line to visualize growth dynamics (Materials and Methods); the initial D10 timepoint was not fit to any condition, as it represents the initial pool of EBs. Colors indicate experimental condition; bold lines indicate the overall growth trend of all batches; thin lines indicate the growth trend of individual batches (biological replicates); single datapoints indicate individual organoids (technical replicates).

### Exogenous ECM exposure influences early organoid morphology and growth dynamics

To assess general features of organoid development, growth, and morphology, we used longitudinal brightfield microscopy imaging (**Fig. 1B**, **Fig. S1**). At D10, EBs had a smooth circular shape and brightening of the outer rim of the tissue, indicating that neuroepithelialisation started (**Fig. 1B**). Morphological changes were rapidly observed in the presence of exECM (**Fig. 1B**, **Fig. S1**; exECM^+D^ and exECM^+L^); organoids appeared irregularly shaped at D13, and tissue budding was visible from D16, most prominently in exECM^+D^ conditions. In the absence of exECM, organoids remained spherical and maintained the outer brightening during the first 20 days (**Fig. 1B**, **Fig. S1**; exECM^-^). Based on these images, organoid diameter was measured (**Fig. 1C**). Growth dynamics were reproducible across different batches of the same cell line within each experimental condition, and largely comparable across all four cell lines. At D10, organoid diameter varied between 600 μm and 800 μm. From D13 to D20, organoid diameter increased more prominently in exECM^+D^, followed by exECM^+L^, and finally exECM^-^ conditions (**Fig. 1C**). Remarkably, evident morphological differences gradually vanished, and organoids appeared identical across conditions and cell lines at D40, with clear tissue budding also in exECM^-^ organoids (**Fig. 1B**, D40). Thus, the presence and concentration of exogenous ECM impacted morphological features and growth dynamics of organoids during neuroepithelium generation and expansion (first 20 days of development) but may not have lasting effects on tissue architecture.

### Exogenous ECM exposure promotes fast rearrangement of neural progenitors

To understand whether exogenous ECM exposure alters neuroepithelial morphology, we assessed early organization of neural progenitors. To visualize the position of individual NPCs within the tissue, we used an H9-derived reporter cell line in which the expression of green fluorescent protein was driven by the SOX2 promoter (SOX2::SOX2-p2A-EGFP, hereafter SOX2::EGFP), marking bona fide neural progenitors (**Sidhaye et al**., 2022). We analyzed organoids containing 80% H9 wild-type (WT) ESCs and 20% H9 SOX2::EGFP ESCs, as this mixing ratio was sparse enough to allow the recognition of individual SOX2::EGFP NPCs, while also revealing their overall tissue distribution. Immunostaining of a member of the atypical protein kinase C subfamily (PKCζ) was used to mark the neuroepithelial apical domain (**Soriano et al**., 2016) (**Fig. 2**).

**Figure 2.**
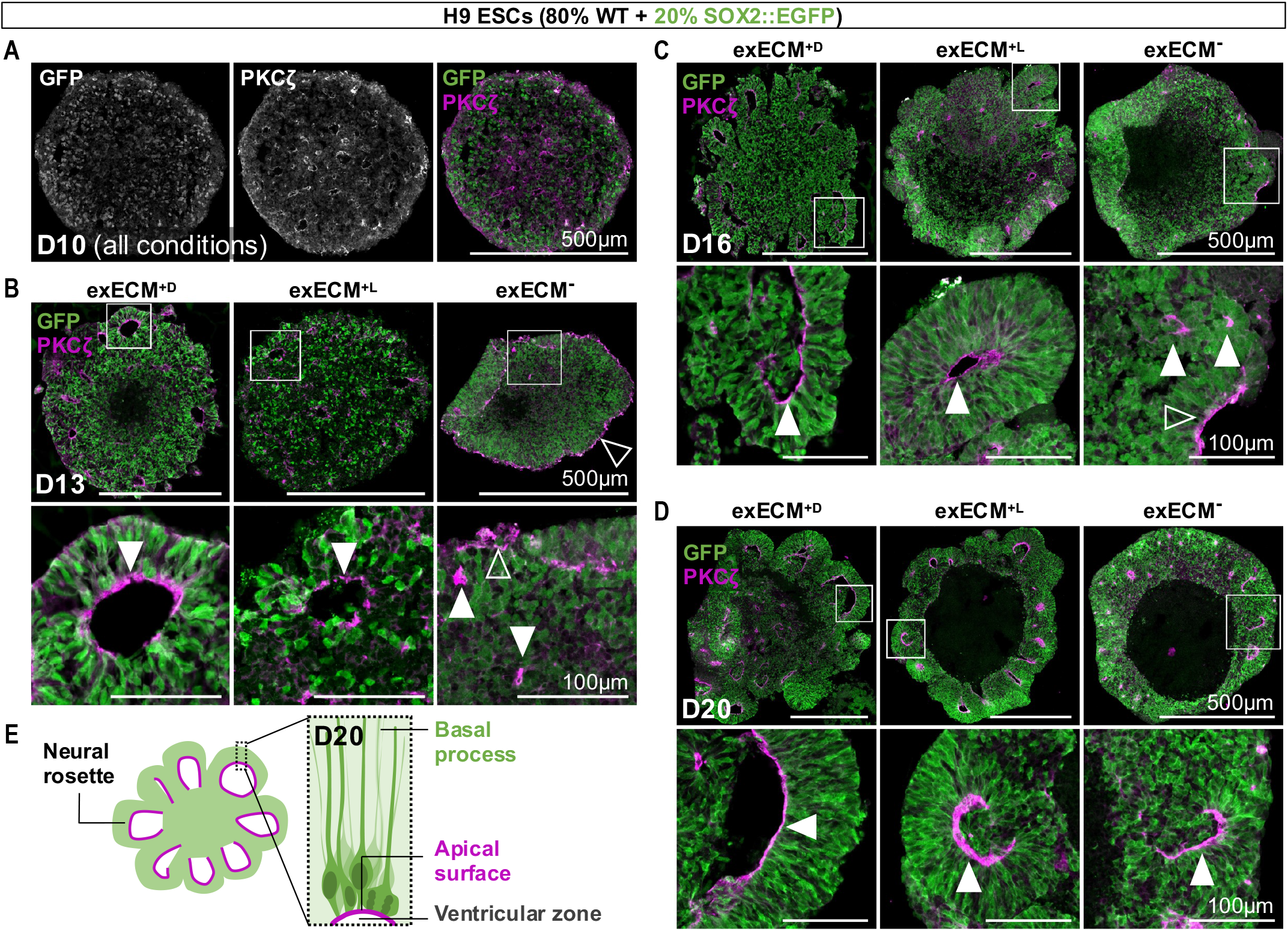
NPCs undergo spatial reorganization and morphological changes during neuroepithelial stages. Organoids generated with 80% WT H9 ESCs and 20% SOX2::EGFP were used to visualize individual neural progenitors at D10 **(A)**, D13 **(B)**, D16 **(C)**, and D20 **(D)**. Bottom panels: magnification of inset. Arrowheads mark the location of PKCζ lining the organoid outer surface (△) or the ventricular zone of neural rosettes (white arrowheads, ▲). **(E)** Schematic representation of the radial organization of neural progenitors in neural rosettes, seen in all conditions at D20.

At D10, small hollow regions - cavitation spots - were present throughout the EB tissue, indicating initial points of structural asymmetry (**Fig. 2A**). However, SOX2::EGFP^+^ neural progenitors appeared scattered, lacking a clear organization; and the direction of apical-basal polarity was yet undefined, with PKCζ staining lining both external and internal surfaces (**Fig. 2A**, **Fig. S2A**). Exposure to exECM led to rapid tissue rearrangements. Within 3 days (**Fig. 2B**, D13), neural progenitors in exECM^+D^ and exECM^+L^organoids were arranged in neural rosettes, acquiring an elongated morphology around large internal lumina (ventricular zones) delimited by a PKCζ^+^ apical surface (**Fig. 2B-D**, ▲; schematized in **Fig. 2E**). In contrast, until D16, exECM-organoids maintained outer apical domains (**Fig. 2B-C**; **Δ**) and few small internal lumina (**Fig. 2B-C**; ▲). At D20, however, neural rosettes with larger lumina and radial progenitor arrangement were also widespread in exECM^-^ organoids (**Fig. 2D**; ▲). Overall, different modes of exogenous ECM exposure caused fast changes in tissue polarity and NPC organization. Interestingly, analogous changes happened in the absence of exogenous ECM with a delay of 5-7 days, suggesting that intrinsic self-organization processes must be in place in exECM^-^ organoids.

### Rosette formation is driven by exogenous or endogenous ECM signaling

To understand what drives the timeline of NPC polarization, we assessed the location of PKCζ and Fibronectin (FN), markers of apical and basal domains, respectively (**Fig. 3A-D**, **Fig. S2–S4**). The FN antibody recognized FN of mouse (Matrigel-derived) and human (endogenously produced) origin. At D10, EBs showed numerous cavitation spots and undefined apical-basal domains (**Fig. S2A**). Embedding in a droplet of exogenous ECM led to the formation of a permanent basal domain on the outer organoid surface, as seen by the surrounding mesh of FN from D13 to D20 (**Fig. 3A-B**, **Fig. S2-4**; exECM^+D^). ExECM dissolution in the culture medium led to the formation of a thin ECM coating at the organoid surface (**Fig. 3A**, **Fig. S2B**, exECM^+L^) that remained visible one week after exECM had been removed (D20, **Fig. 3B**, **Fig. S3–4**). Remarkably, exECM^-^ organoids showed abundant endogenous FN production from early developmental stages. At D13 (**Fig. 3A**, **Fig. S2B**) and D16 (**Fig. S3**), FN was mostly scattered across the tissue of exECM^-^ organoids, whereas, at D20, an apical-basal axis was established, with circular arrangement of FN^+^ regions around PKCζ^+^ lumina of neural rosettes (**Fig. 3B**, **Fig. S4**). In summary, exogenous ECM addition established a clear basal-out/apical-in polarity axis in exECM^+^ organoids and drove rosette formation from D13; and exECM^-^ organoids endogenously produced ECM components that similarly culminated in rosette arrangement by self-organization (schematized in **Fig. 3C-D**).

**Figure 3.**
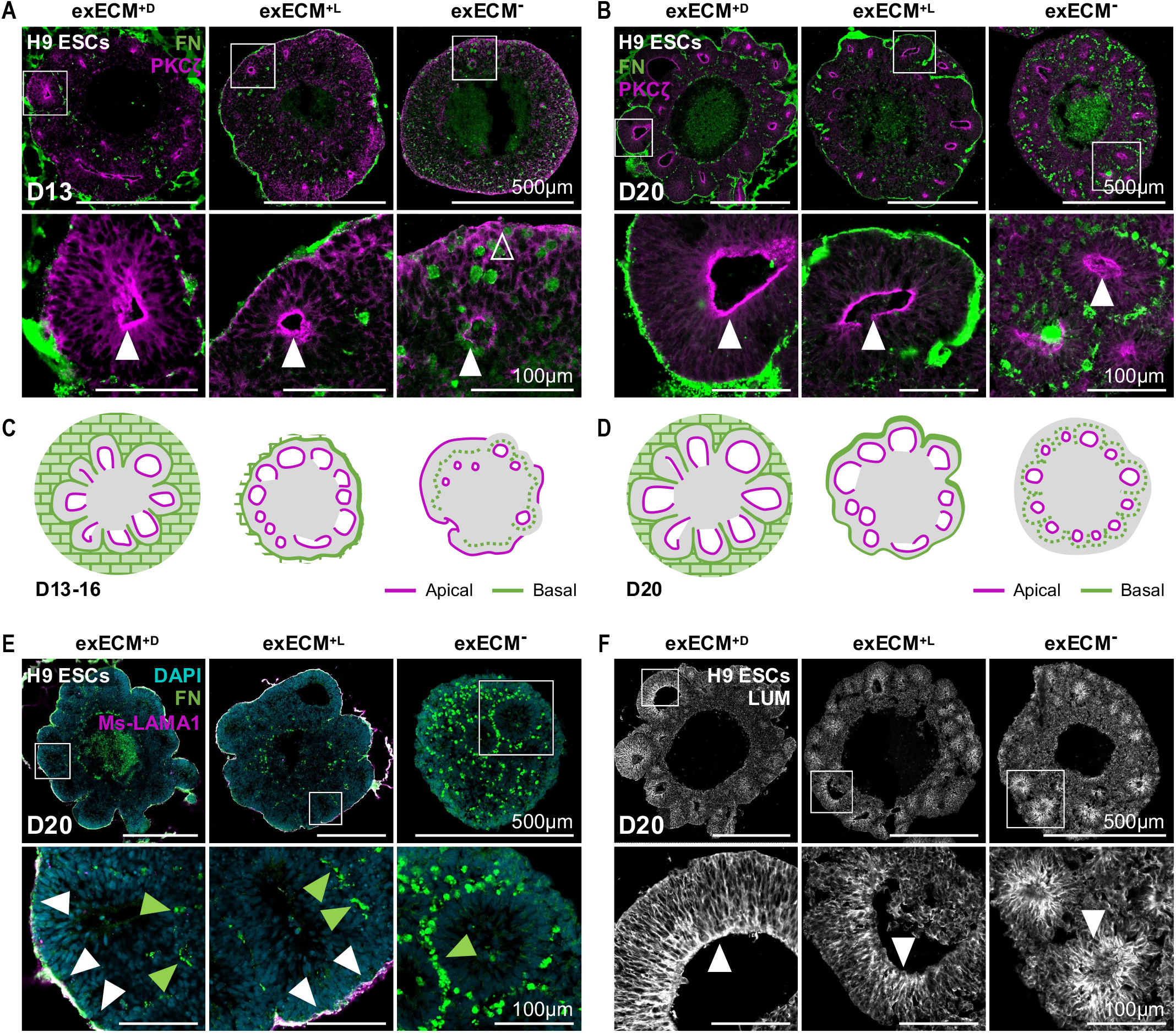
Exogenous and endogenous ECM proteins participate in the establishment of apical-basal polarity. **(A-D)** Apical and basal domains are marked by PKCζ and FN, respectively; arrowheads mark the location of PKCζ lining the organoid outer surface (Δ) or the ventricular zone of neural rosettes within the tissue (▲). An apical-basal polarity axis is defined at D13 in exECM^+D^ and exECM^+L^ organoids **(A)** and at D20 in exECM^-^ organoids **(B)**(schematic representations in. **(E)** Ms-LAMA1 antibody can be used to identify mouse-derived ECM (Matrigel). At D20, exECM^+D^ and exECM^+L^ organoids show a coating of Ms-LAMA1 originating from Matrigel, which co-localizes with FN (▲); also, FN^+^ but Ms-LAMA1^-^ speckles are seen within the tissue, indicating endogenously-produced FN (green arrowheads, 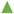). ExECM^-^ organoids show abundant endogenously-produced FN surrounding neural rosettes 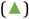. **(F)** Lumican is abundantly produced in all experimental conditions and localizes to the epithelial apical surface, lining the ventricular zone of neural rosettes (▲). Bottom panels: magnification of inset.

Interestingly, the patterns of FN^+^ regions at D20 were remarkably different in exECM^-^ and exECM^+^ conditions (**Fig. 3B and D**). This prompted us to discriminate between ECM produced endogenously and ECM contributed by Matrigel, using an antibody that recognizes mouse, but not human, laminin-α1 (Ms-LAMA1) (**Fig. 3E**). In exECM^-^ organoids, Ms-LAMA1 was absent, as expected; FN showed a speckled pattern around neural rosettes and did not reach the outer-most surface of the organoids - a pattern indicative of endogenously-produced ECM. In exECM^+^ organoids, however, the smooth FN^+^ surface was co-positive for Ms-LAMA1, identifying Matrigel-derived ECM. In addition, FN-positive but Ms-LAMA1-negative speckles were seen within the tissue in exECM^+^ organoids, indicative of endogenous FN production. Thus, Matrigel addition led to the formation of a sheet of ECM at the outer-most organoid surface, distinguishable from, but not replacing, endogenously-produced ECM within the tissue.

Several ECM proteins present in the developing brain *in vivo* are absent from Matrigel. Finally, we assessed the presence and tissue distribution of one such ECM component, Lumican (LUM), which is produced by human NPCs and plays an important role in cortical development (**Long et al**., 2018). LUM was abundant in organoids from early stages of development, and its tissue distribution followed a pattern similar to that of PKCζ. Specifically, we observed scattered and disordered distribution of LUM at D10 (**Fig. S5A**) and accumulation of LUM in rosette lumina from D13 in exECM^+^ organoids (**Fig. S5B**), and at D20 in all conditions (**Fig. 3F**, **Fig. S5C**). These observations show that Matrigel exposure does not influence the endogenous generation of other ECM components. Furthermore, it reiterates that both exogenously-guided and self-guided tissue organization culminate in comparable polarized arrangements at D20 of culture, along with the formation of neural rosettes.

### Rosette architecture and identity are comparable at early neurogenic stages

To assess if the initial differences in neuroepithelisation impact the relative organization of neural progenitors and neurons, we evaluated organoids at neurogenic stages. Independently of exposure to exogenous ECM, the first neurons (MAP2^+^) appeared at around D20 of organoid development, typically on the outside of neural rosettes (Fig. S6). We re-evaluated tissue architecture at D40, when neurogenesis is in a more advanced phase and prominent neural rosettes were visible in all conditions with brightfield imaging (Fig. 1B, Fig. S1, tissue architecture schematized in Fig. 4A). At this stage, rosettes maintained a well-defined apical-basal axis (Fig. S7) and SOX2^+^ neural progenitors were radially arranged around large lumina and surrounded by abundant MAP2^+^ neurons (Fig. 4B, Fig. S8). Also, rosettes presented a dorsal-cortical identity and a stereotypical inside-out organization of radial glia (SOX2^+^), intermediate progenitor cells (TBR2^+^), and early-born excitatory neurons (CTIP2^+^) (Fig. 4C, Fig. S9, schematized in Fig. 4A’). Small clusters of DLX2^+^ interneuron progenitors were only occasionally seen (Fig. S10A), which is characteristic of a predominantly dorsal fate. Measurement of organoid diameter (Fig. 4E) and median rosette area (Fig. 4F) showed that exECM^+D^ organoids were significantly larger and presented larger rosette area than exECM^-^, whereas differences between exECM^+L^ and exECM^-^ were less- or non-significant. Thus, during production of deep-layer excitatory neurons, general features of polarity maintenance, cellular organization, and tissue architecture were largely independent of early exECM exposure or hPSC genetic background.

**Figure 4.**
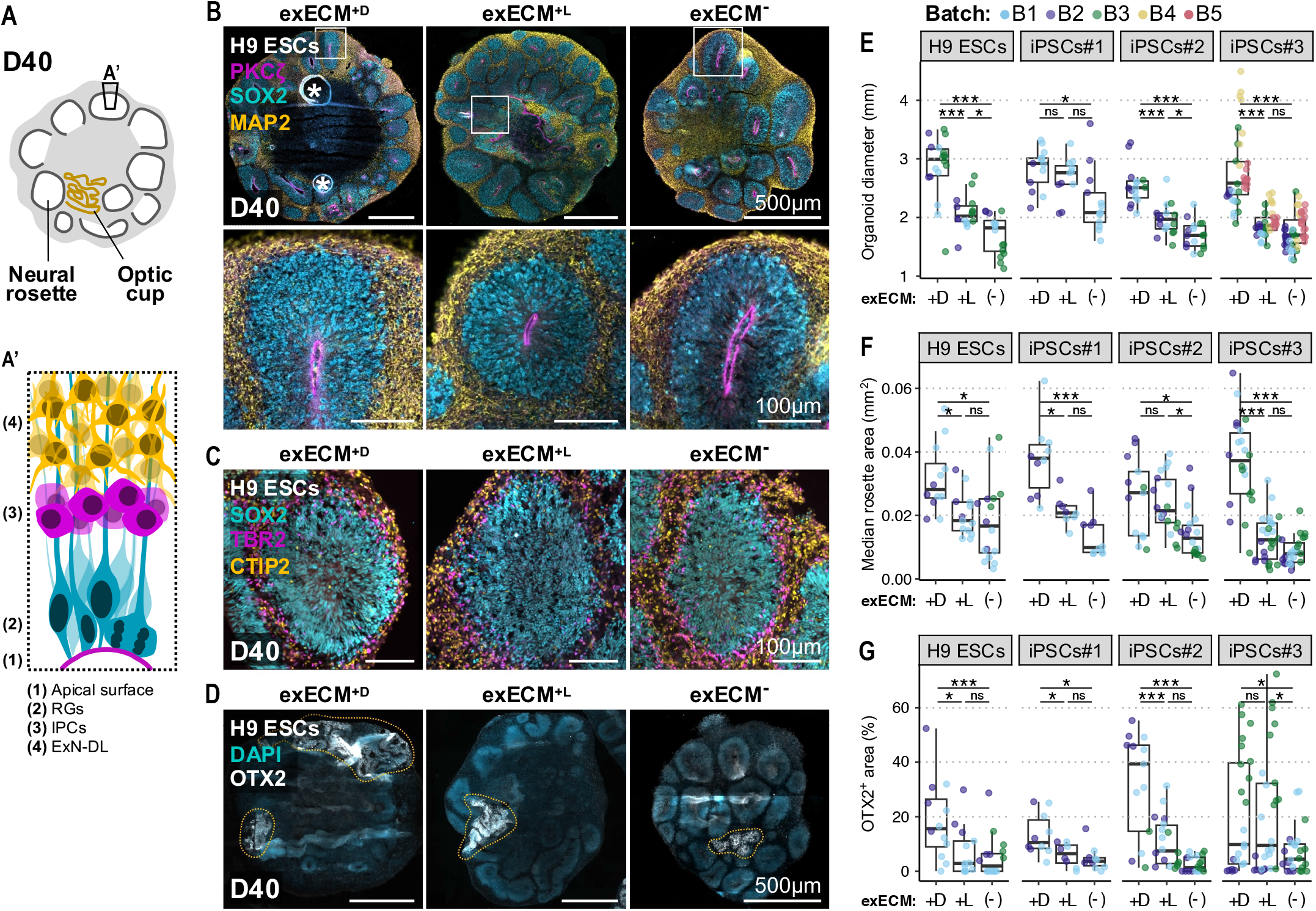
At D40, the organoid tissue is mostly comprised of dorsal-cortical neural rosettes and minor regions of mis-patterned cells. (A) Schematic representation of tissue architecture at D40. Abundant neural rosettes and smaller regions of optic cup tissue are present. (A’) Schematic representation of dorsal-cortical rosette organization. (B) Organoids from all conditions show abundant neural rosettes with PKCζ^+^ ventricular zone, and inside-out organization of SOX2^+^ neural progenitors and MAP2^+^ neurons. Bottom panels: magnification of inset; *: staining artifact. (C) Neural rosettes have dorsal-cortical identity, and radial glia (RGs, SOX2^+^), dorsal intermediate progenitors (IPCs, TBR2^+^) and early-born deep-layer excitatory neurons (ExN-DL, CTIP2^+^) show layered arrangement. (D) Non-telencephalic tissue shows convoluted and disorganized morphology and is marked by OTX2 (and TTR - see Fig. S10B; demarcated by dashes). Organoid diameter (E), median rosette area (F), and percentage of OTX2+ area (G) across experimental conditions and cell lines. Each datapoint is an individual organoid (technical replicate), datapoint colors indicate organoid batches (biological replicates). Ordinary one-way analysis of variance (ANOVA); 0 ≤ p < 0.001, ***; 0.001 ≤ p < 0.01, **; 0.01 ≤ p < 0.05, *; p ≥ 0.05, ns (see results of statistical tests in ibles S1-S3).

### Exogenous ECM permanence in the organoid tissue contributes to mispatterning

Tissue mis-patterning due to protocol variability or undesired exogenous cues can cause the presence of unwanted regional fates within organoids. While examining organoid morphology at D40, we identified regions that did not organize in neural rosettes, appearing more convoluted and disordered (**Fig. 4D**, schematized in **Fig. 4A**). Co-expression of OTX2 and TTR indicated optic-cup identity (**Fig. 4D**, **Fig. S10B**). By quantifying the percentage of OTX2^+^ tissue area per organoid, we found that although the extent of OTX2^+^ regions was batch-dependent, it was significantly higher in exECM^+D^ organoids in all cell lines and had a trend to be higher in exECM^+L^ than in exECM^-^ organoids, which was only significant in iPSCs#3-derived organoids (**Fig. 4F**). Importantly, Ms-LAMA1 staining showed that most exECM^+D^ organoids remained encapsulated in a Matrigel droplet, while remnants of Matrigel were still visible within exECM^+L^ organoids at this stage (**Fig. S11A**). Overall, organoids cultured in the absence of exogenous ECM were more homogenous, and continued signaling from exogenous ECM potentiated an increased differentiation or expansion of non-telencephalic tissue.

### Long-term organoid maturation is independent of early exECM exposure

One of the main goals of organoid modeling is to gain access to mature features of developing neuronal tissue. Thus, to evaluate potential long-term effects of differential early exposure to exogenous ECM on organoid maturation, we investigated the cellular composition of organoids at a developmental stage when different classes of mature neurons are present (D120). Strikingly, although the organoid tissue had by this time completely outgrown the exECM added early on, Matrigel remained visible as large solid formations in exECM^+D^ organoids, or as small remnants in exECM^+L^ organoids (**Fig. S11B**). To perform an unbiased analysis of cell-type composition, we resorted to single-cell RNA sequencing (scRNAseq), focusing on H9-derived organoids and three organoids per condition (**Fig. 5A-H, Fig. S12A**). Although exECM^-^ organoids were smaller than exECM^+L^ and exECM^+D^ organoids (**Fig. 5A;** also observed in other cell lines: **Fig. S13A**), no other relevant morphological distinctions were found. To preserve organoid origin information for each cell, we used tagging with a unique molecular identifier oligonucleotide and computational demultiplexing after sequencing; the majority of cells recovered had unique barcodes, yielding 14.5k high quality cells used for further analysis (**Materials and Methods**, **Fig. S12A-C**).

**Figure 5.**
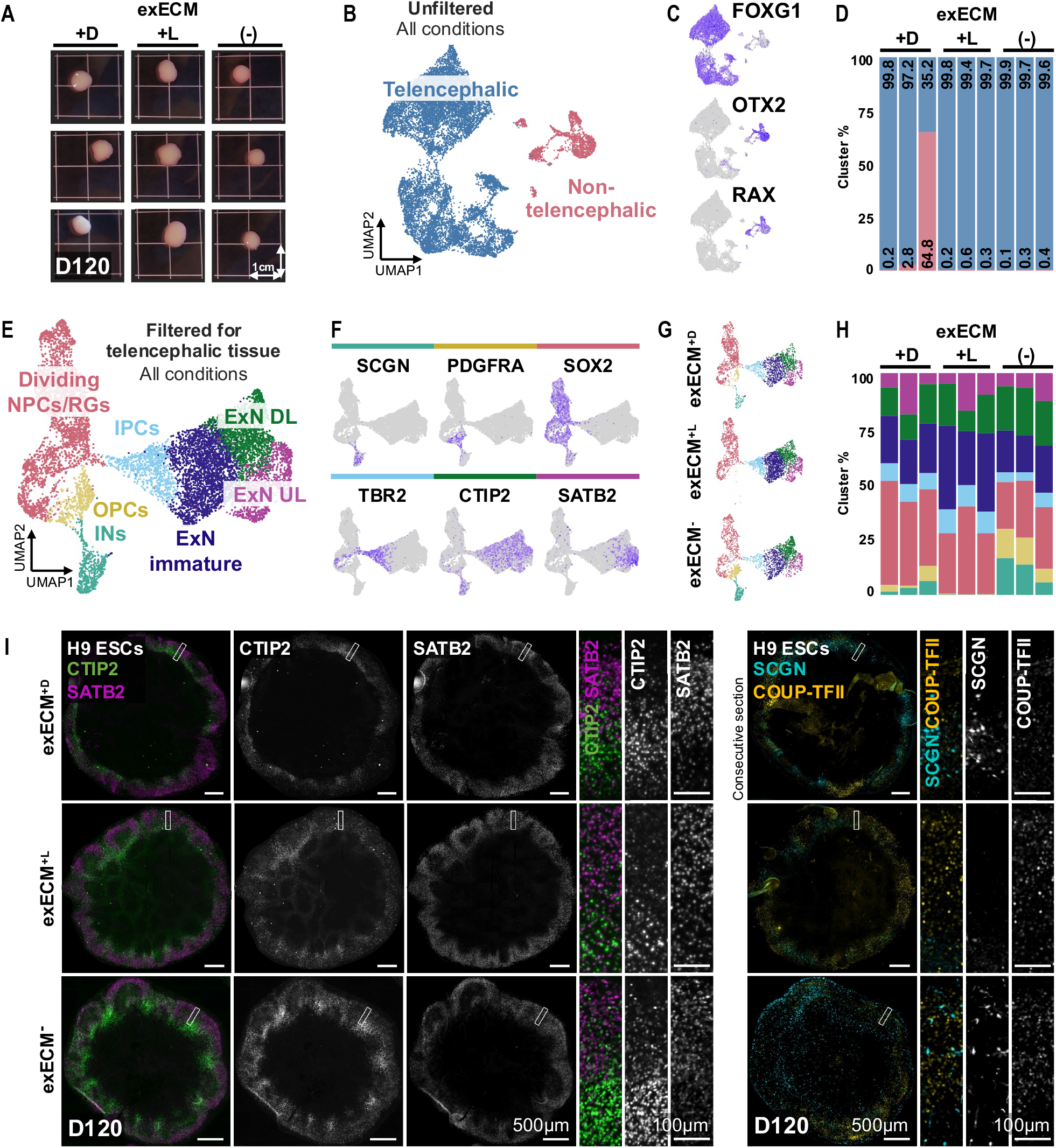
Mature organoid culture produces comparable cell types across conditions. **(A)** At D120, nine H9 ESCs-derived organoids were used for scRNAseq analysis. **(B-C)** UMAP projection of cells isolated from each organoid identifies two main clusters: FOXG1^+^ telencephalic cells; and OTX2^+^RAX^+^ non-telencephalic cells. **(D)** Non-telencephalic cells derive mainly from one exECM^+D^ organoid, making up over 60% of the tissue. **(E)** UMAP projection after filtering and exclusion of non-telencephalic clusters allows the identification of 8 clusters, including dividing NPCs/radial glia progenitors (RGs), oligodendrocyte precursor cells (OPCs), interneurons (INs), intermediate progenitor cells (IPCs), immature excitatory neurons (ExN), deep-layer excitatory neurons (ExNs DL) and upper-layer ExNs (ExNs UL), marked by cell-type specific marker genes **(F)**. **(G-H)** Calculation of the percentage of cells per cluster per organoid shows an increase in INs and OPCs in exECM^-^ organoids. **(I)** Tissue immunostaining of organoids at D120 shows abundant deep- and upper-layer neurons (CTIP2^+^ and SATB2^+^, respectively) with rudimentary layer organization, as well as less abundant populations of interneurons (SCGN^+^and COUP-TFII^+^).

Unsupervised clustering in UMAP projection identified two main clusters (**Fig. 5B**). Differential gene expression analysis revealed that the most abundant cluster comprised FOXG1^+^ telencephalic cells, whereas the other cluster was positive for non-telencephalic markers, including OTX2 and RAX (**Fig. 5C**) and originated mostly from a single exECM^+D^ organoid (**Fig. 5D**). Thus, the second cluster represented a mis-differentiation to optic cup that was already predominant in exECM^+D^ organoids at D40 (**Fig. 4D and G**). After filtering out low-quality cells and removing non-telencephalic clusters, we separated clusters of dividing NPCs and radial glia progenitors (RGs), oligodendrocyte precursor cells (OPCs), interneurons (INs), intermediate progenitor cells (IPCs), immature excitatory neurons (ExNs), deep-layer ExNs (ExNs DL) and upper-layer ExNs (ExNs UL) (**Materials and Methods** and **Fig. 5E**), all marked by expression of cell-type specific marker genes (**Fig. 5F**, **Fig. S12D**). Calculation of the percentage of cells per cluster indicated that exECM^-^ organoids contained more ventral progenitors and interneurons (**Fig. 5G-H**). However, the proportions of cell types along the excitatory lineage (excluding OPCs and INs) were comparable between conditions (**Fig. S12E-G**). This suggested that slight patterning differences along the dorsal-ventral axis of the telencephalon may be introduced by differential exECM exposure and sustained after long-term culture.

To validate the cell-type composition in all cell lines and conditions, we resorted to immunostaining (**Fig. 5I**, **Fig. S13B**). Organoids were composed mostly of deep- and upper-layer ExNs (CTIP2^+^ and SATB2^+^, respectively) with rudimentary layer organization, a finding that matched the tissue identity verified at D40 (**Fig. 4B**, **Fig. S9**, **Fig. S10A**), the indications from the scRNAseq data, and the fact that a guided differentiation protocol was used. A small number of interneurons (SCGN^+^ and COUPTFII^+^, indicating caudal ganglionic eminence origin) was found intermingled with ExNs in all conditions and cell lines (**Fig. 5I**, **Fig. S13B**). Overall, although the exact proportions of cell types showed slight variations, global cell-type composition and developmental stage were highly similar across experimental conditions at D120, indicating that exposure to exogenous ECM at the neuroepithelial stage holds few long-lasting effects in organoid development.

## Discussion

Epithelial morphogenesis involves the consecutive coordination of several processes, including tissue polarization by cell-ECM and cell-cell interactions, and lumen formation, maintenance, and expansion (**Datta et al**., 2011; **Martín-Belmonte & Mostov**, 2008). Given its central role in defining tissue morphology, a critical component of epithelial systems is the source of ECM.

### Mechanisms of tissue morphogenesis

Here, we characterized how the presence or absence of exogenous ECM affects the development of human telencephalic organoids. We found that, in the absence of exECM, early spots of cavitation initiate tissue asymmetry and are likely the starting points of rosette lumina. These observations suggest a mechanism resembling secondary neurulation *in vivo*, which has also been reported in 2D neural progenitors (**Fedorova et al**., 2019; **Hříbková et al**., 2018) and in other *in vitro* epithelial systems, such as Madin-Darby canine kidney (MDCK) cysts (**Martín-Belmonte et al**., 2008; **Yu et al**., 2005). Furthermore, in agreement with observations made in early studies with embryoid bodies (**S. Li et al., 2002**, **2003**), exECM^-^ organoids produce ECM proteins - such as Fibronectin and Lumican - that self-assemble and define an apical-basal polarity axis at pre-neurogenic stages. This suggests that ECM-producing cells are neural progenitors, which is in line with a study showing that organoid radial glia cells express ECM components comparable to those produced by the endogenous population (**Camp et al**., 2015). In the human brain, NPCs have been postulated to contribute to basal deposition of ECM constituents via vesicular transport in their basal processes (**Fietz et al**., 2012), thereby polarizing their own niche environment. We propose that an analogous self-sustained process may be taking place during *in vitro* development, contributing to the establishment and maintenance of apical-basal polarity in the absence of exogenous instructive signals.

Although polarization and lumen formation emerge spontaneously *in vitro*, both processes can be influenced by culture conditions. Here, the action of exogenous ECM is likely two-fold: 1) introduction of a strong basement membrane signal at the organoid surface; and 2) signal amplification by recruitment and polymerization of endogenously-produced ECM. In fact, Fibronectin staining within the organoid tissue at D13-16 is widespread in exECM^-^ organoids and sparser in exECM^+^ organoids; these differences may be due to recruitment and assembly of endogenously-produced FN at the organoid surface in the presence of exECM. Laminin likely plays a role in these processes, as it constitutes around 60% of Matrigel (**Corning Incorporated Life Sciences**, 2016), and has been shown to form the initial cell-anchored polymer needed for subsequent ECM assembly, and to nucleate the polymerization of other ECM proteins (**Cheng et al**., 1997; **S. Li et al., 2002, 2003**). Thus, slow assembly of endogenous ECM is overtaken by a mass-action of exogenous ECM upon Matrigel exposure, leading to a quick polarization process that likely occurs through a different molecular mechanism than that seen in exECM^-^ organoids. As such, exECM supplementation can be experimentally advantageous to assess NPC arrangement and morphology with short experimental timelines, as has been applied in evolutionary (**Benito-Kwiecinski et al**., 2021) and disease modeling studies (**Krenn et al**., 2021).

Another important conclusion is that Matrigel exposure in the form of a jellified matrix (exECM^+D^) or transient dissolution in the medium (exECM^+L^) does not critically affect early tissue morphogenesis. In fact, both formulations result in the accumulation of ECM proteins at the surface of the organoids, leading to polarity establishment and rosette formation within 3 days. This is in contrast with other organoid systems in which both exECM presence and jellification are needed to establish apical-basal polarity (**Inman & Bissell**, 2010; **Kakni et al**., 2022; **Plachot et al**., 2009). Furthermore, early morphological rearrangements are maintained in exECM^+L^ organoids after Matrigel is removed from the culture medium, indicating that continuity of exposure is also unnecessary. Contrarywise, reversal of apical-basal polarity by withdrawal of exogenous ECM proteins from culture has been extensively applied in other organoid models to facilitate access to the apical epithelial surface (**Co et al., 2019**, **2021**; **Krüger et al**., 2020; **Y. Li et al**., 2020; **Nash et al**., 2021; **Salahudeen et al**., 2020; **Stroulios et al**., 2021). Overall, in other systems, morphogenic rearrangements require a high concentration of exogenous ECM and/or the physical support and constraint provided by a jellified matrix; as well as continuity an exogenous ECM source to maintain polarity. All these are dispensable in the human telencephalic organoids described here, possibly because of the self-sustainable ECM production within the organoid tissue.

The impact of endogenous and exogenous ECM sources on epithelial morphogenesis has been addressed in other *in vitro* systems. An example are experimental paradigms that show the formation of somites, for which either a low concentration of Matrigel in the culture medium (**Sanaki-Matsumiya et al**., 2022; **van den Brink et al**., 2020; **Veenvliet et al**., 2020) or differentiation of ECM-producing cells within the tissue are required (**Amadei et al**., 2022; **Bao et al**., 2022; **Lau et al**., 2022). Similarly, breast luminal epithelial cells can be polarized *in vitro* by exogenous addition of laminin-1, or co-culture with ECM-producing myoepithelial cells (**Gudjonsson et al**., 2002). Our findings indicate that although polarization and lumen formation of neuroepithelial cells can be rapidly potentiated by exogenous ECM exposure, the presence of ECM-producing tissue-resident cells is enough to intrinsically drive self-organization. Ultimately, exogenous ECM addition and ECM-producing cells offer alternative paths for reaching comparable morphogenic outcomes

### Organoid development at neurogenic stages

From the peak of production of deep-layer neurons (D40) to stages reminiscent of late corticogenesis (D120), organoid development proceeds in analogous ways independently of early exposure to exogenous ECM. In all conditions and hPSC genetic backgrounds, the tissue is organized in neural rosettes with comparable dorsal-cortical identity, inside-out organization of progenitors and neurons, and apical-basal polarity axis. NPCs give rise to, first, CTIP2^+^ deep-layer neurons and, later, SATB2^+^ upper-layer neurons. Transcriptional features and rudimentary layering of mature cortical neurons are acquired equally across conditions, as seen by scRNAseq and tissue staining, respectively. The populations of interneurons identified with these assays may be derived from small regions of ventrally-patterned tissue seen early stages, or a product of the differentiation of cortical progenitors into both excitatory and inhibitory neurons (**Delgado et al**., 2021). Despite these commonalities, biases in tissue patterning that favor the expansion of optic cup tissue in exECM^+D^ organoids are already significantly higher at D40 and persist throughout time. This suggests a dependency on continuity of exposure, concentration or physical action of exogenous ECM, which is in agreement with pioneering studies on *in vitro* differentiation of the optic cup (**Eiraku et al**., 2011). Taken together, these data show that general features of long-term maintenance and maturation of organoids are independent of early exposure to exogenous ECM. These conclusions are corroborated by organoid studies performed without exECM exposure, in which features of neuronal maturation, such as migratory patterns and electrophysiological activity, were successfully achieved (**Birey et al**., 2017; **Paşca et al**., 2011).

### Limitations of the study

Here, we explored the effect of exposure of dorsal telencephalic brain organoids to an exogenous ECM preparation, Matrigel, at the stage of neuroepithelium formation. We provide a detailed morphological analysis of the organoid tissue at early stages, mainly focusing on apical-basal polarity and ECM composition, but the exact molecular mechanisms of rosette assembly were not explored. In the future, it will be interesting to perform a more detailed biochemical characterization of cavitation and polarization processes, coupled with live imaging of mosaic reporter PSC-derived organoids at early stages, to visualize tissue morphogenesis. Additionally, we chose to use Matrigel, it being the golden-standard exogenous ECM in the cerebral organoid field. However, Matrigel contains an incompletely defined mix of ECM components and growth factors, which poses challenges in the interpretation of experimental results. In fact, although we propose a likely action of Matrigel-derived Laminin in polarity establishment, we are unable to precisely pinpoint the component or components responsible for the observed effects in exposed cultures. For example, tissue patterning effects observed in exECM^+^ organoids may be attributable to unknown growth factors introduced in the culture, which would be impossible to track and control for, given the batch-to-batch variability of Matrigel. Moving forward, testing the action of purified ECM components in analogous experimental paradigms would certainly help uncover these questions. These would also follow recent attempts to tackle the variable and undefined composition of Matrigel by developing synthetic alternatives and Matrigel-free culturing methods (**Aisenbrey & Murphy**, 2020; **S. Kim et al**., 2022; **Kozlowski et al**., 2021; **Kratochvil et al**., 2019; **Nayler et al**., 2021; **Roth et al**., 2021). With our study, we provide a robust, reproducible, and scalable assay that can be used to explore these hypotheses. Finally, we focused on the action of exogenous ECM at specific stages of neuroepithelium formation and morphogenesis. Certain protocols suggest that addition at later timepoints or throughout long-term organoid culture may improve cortical plate formation (**Bhaduri et al**., 2020; **Kadoshima et al**., 2013; **Lancaster et al**., 2017; **Velasco et al**., 2019), the implications of which were out of the scope of this study.

### Impact and future directions

Supplementation of organoid cultures with exogenous ECM is widespread, but a full characterization of its necessity and impact is missing for certain *in vitro* systems, including cerebral organoids. In summary, our data strongly support an experimental model in which early exogenous ECM exposure is useful to trigger quick morphogenic changes, when experimentally needed, but not necessary for long-term development of human telencephalic organoids. Key findings are supported by a comparable study carried out in cerebellar organoids (**Nayler et al**., 2021), showing how they may be transversal to neuroectoderm-derived tissues and applicable to *in vitro* models of other brain regions. The implications of these conclusions are manyfold. On the one hand, they generate crucial knowledge regarding neuroepithelial biology *in vitro* and the self-sustainability of tissue morphogenesis without the need for extrinsic guiding factors. On the other hand, they call for a re-evaluation of the necessity to use exogenous ECM preparations in neuroepithelial model systems. In fact, exposure to mixed or pure ECM components, albeit often useful and essential, are brute-force approaches to recapitulate endogenous processes. Thus, culture components that introduce strong signaling in epithelial tissues should be used mindfully and the necessity and consequences of their presence considered. Ultimately, adding to previous efforts from other groups, we believe that this study answered long-standing questions in the brain organoid field and is a positive step towards fully characterizing and unifying brain organoid research.

## Data and code availability

The scRNAseq data discussed in this publication have been deposited in NCBI’s Gene Expression Omnibus (**Edgar et al**., 2002) and are accessible through GEO Series accession number GSE220085. Analysis was performed as outlined in the **Materials and Methods**. No custom code was generated for this study; used code is available upon request.

## Acknowledgements

We thank Oliver Eichmüller for help with analyses, and members of the Knoblich lab for feedback on the manuscript. We thank the IMBA stem cell core facility for cell reprogramming services; the IMBA/IMP/GMI BioOptics facility for microscopy services; and the IMBA/IMP/GMI Bioinformatics for sequencing analysis. Work in the Knoblich laboratory is supported by the Austrian Academy of Sciences, the Austrian Science Fund (FWF), (Special Research Programme F7804-B and Stand-Alone grants P 35680 and P 35369), the Austrian Federal Ministry of Education, Science and Research, the City of Vienna, and a European Research Council (ERC) Advanced Grant under the European Union’s Horizon 2020 programs (no. 695642 and no. 874769). JS was supported by the EMBO long term fellowship (EMBO ALTF 794-2018) and the European Union’s Horizon 2020 research and innovation program under the Marie Skłodowska-Curie fellowship agreement 841940.

## Author contributions

Conceptualization: CMC, NSC, JS, JAK; Methodology: CMC, VP, AP, PM; Formal analysis and investigation: CMC; Writing - original draft preparation: CMC; Writing - review and editing: CMC, NSC, JS, JAK; Funding acquisition: JAK; Resources: JAK; Supervision: NSC, JS, JAK.

## Materials and methods

### hPSC maintenance and passaging

Feeder-free hESC line WA09 (H9 ESCs) were commercially obtained from WiCell; iPSCs#1 and iPSCs#2 were reprogrammed in-house from fibroblasts of healthy donors (internal nomenclature: iPSCs 178/4 and iPSCs 176/1, respectively); iPSCs#3 were Rozh-5 iPSCs commercially obtained from HipSci. Therefore, we used a very common ESC line, as well as commercially-available and in-house reprogrammed iPSC lines. All cells were cultured on 6 well-plates (Corning, 3516) coated with hESCs-qualified Matrigel (Corning, 354277), and maintained in complete mTeSR1 medium (StemCell Technologies, 85875). Cells were fed daily with 2 mL of mTeSR1 (feeding with 4 mL allowed skipping of one feeding day per week) and passaged after reaching 60-80% confluency (every 3-5 days). For passaging, cells were exposed to 0.5 mM EDTA diluted in PBS (pH 7.4, without MgCl_2_ or CaCl_2_; Gibco, 14190-250) during 3 min at 37 °C, lifted in mTeSR1 by gentle spraying of the bottom of the well, and triturated to small clusters of 20-50 cells. Cells were routinely tested for mycoplasma, and verified to display a normal karyotype resorting to short tandem repeat (STR) analysis and intact genomic integrity resorting to single-nucleotide polymorphism (SNP) analysis. Cells, embryoid bodies (EBs), and organoids were kept in a 5 % CO_2_ incubator at 37 °C.

### Dorsal tissue-enriched telencephalic organoid generation

Media formulations can be found in **Table S4**. Feeding volumes in 96WP format were of 150 μL, and in 10 cm dish format of 15 mL.

#### Embryoid body formation

Dorsal forebrain-enriched telencephalic organoids were generated as previously described with slight modifications (**Esk et al**., 2020). hPSCs were grown to 60-80% confluency and dissociated into a single cell suspension by Accutase (Sigma-Aldrich, A6964) treatment for 5 min at 37 ºC, followed by manual trituration. Cells were seeded in an ultra-low binding 96-well plate (Szabo-Scandic, COR7007), at a density of 9000 live cells/well, in 150 μL of complete Essential 8 (E8) medium (Thermo Fisher Scientific, A1517001) with 50 μM Rho-associated protein kinase (ROCK) inhibitor (Selleck Chemicals, S1049). On day 3, the medium was replaced with E8 without ROCK inhibitor supplementation. From day 6, EBs were fed daily with Neural Induction (NI, **Table S4**) medium.

#### Batch quality control assessment

On Day 10, batches in which over 80% of EBs formed successfully were kept for further experiments. Quality criteria included EB size above 500 μm, round morphology, and the appearance peripheral tissue clearing, indicative of the start of neuroepithelium formation. Batches compliant with these criteria were randomly divided at D10 into three groups of different conditions of exogenous ECM (exECM) supplementation. Details of handing for each condition described below. Of note, within successful batches, organoid quality was comparable across all conditions and, thus, likely determined by factors preceding or independent from MG exposure, such as pluripotency state, confluency, and passage number of the starting population of hPSCs, as well as user technique, at the stage of EB setup.

#### Tissue patterning

On day 13, the NI medium was replaced with Differentiation Medium without vitamin A (Diff-A, **Table S4**). Two pulse applications of 3 μM of GSK-3 Inhibitor CHIR99021 (Merck Millipore, 361571) were done on days 13 and 14; the medium was replaced with Diff-A without CHIR on day 16.

#### Long-term culture

On day 20, all organoids, regardless of Matrigel condition, were transferred to 10 cm dishes and cultured on an orbital shaker (Celltron) at a rotating speed of 57 rpm. On day 25, the medium was replaced with Differentiation Medium with vitamin A (Diff+A, **Table S4**). From day 25 onwards, the medium formulation remained unaltered throughout organoid development, and the medium was changed twice a week (every 3-4 days).

### Exogenous ECM application

When used, exogenous ECM was Matrigel LDEV-Free (Corning, 354234).

#### No exECM (exECM^-^)

At D10, EBs remained in 96-well plates and received daily exchange of medium according to the schedule described above. Exceptions occurred during weekends, when often one day of feeding was skipped; and during the pulse application of CHIR99021, when medium exchange occurred precisely on D13 and D14 and not on D16.

#### Droplet embedding (exECM^+D^)

At D10, EBs were embedded into droplets of Matrigel on dimpled sheets of parafilm, as previously described (**Lancaster et al**., 2017). Only organoids that remained within exECM droplets were used for downstream analyses; organoids that detached from the droplet were discarded.

#### Liquid embedding (exECM^+L^)

The culture medium (NI) was supplemented with Matrigel at a concentration of 2%V/V at D10. Matrigel was left out from the following feeding, at D13.

Until D20, exECM^+D^ and exECM^+L^ organoids were cultured in suspension, in a stationary 10 cm dish, with media exchange every 3 days, according to the schedule of media described above. Exceptions occurred during the pulse application of CHIR99021, when medium exchange occurred precisely on D13 and D14 and skipped on D16. To avoid organoid attachment, 10 cm dishes were coated with Anti-adherence rinsing solution (Stemcell technologies, 7010) during 2 min, followed by one rinse with PBS, immediately before organoid transfer.

### Cryosectioning

Organoids were collected at days 10, 13, 16, 20, 40, and 120, and fixed in 4% paraformaldehyde (PFA) for 30min (D10-D20) or 1-2 hours (D40D120) at room temperature (RT). After three 15 min washes with PBS, organoids were immersed in a 15% Sucrose (Merck Millipore, 84097)/10% Gelatin (Sigma, G1890-500G) solution at 37 °C, until they sunk to the bottom of the tube (from 1h up to overnight). Organoids were embedded in the same sucrose/gelatin solution, solidified for 30min at 4 ºC, and flash frozen in a bath of 2-methylbutane super-cooled to a temperature of −50 °C by dry ice. Samples were stored at −70 ºC until further processing. Cryoblocks were sectioned at 20 μm thickness using a cryostat (Thermo Fisher Scientific, CryoStar NX70).

### Immunohistological staining

After the slides were defrosted and hydrated during 5 min in PBS, cryosections were permeabilized and blocked with blocking solution (5% bovine serum albumin (BSA; Europa Bioproducts, EQBAH-0500) and 0.3% Triton X-100 (Merck Millipore, 93420) in PBS) for 1-2 hours at RT. Antibody incubations were done in antibody solution (1% BSA, 0.1% Triton X-100 in PBS); after dilution of antibodies, the solution was spun at maximum speed for 2 min. First, sections were incubated with antibody solution containing 1:200 diluted primary antibodies, during 3.5 hours (up to overnight) at RT. Then, after one wash of 5 min with PBS, sections were incubated with antibody solution containing 1:500 diluted secondary antibodies and 1:10,000 diluted Hoechst 33342 nuclear dye (Thermo Fisher Scientific, H3569), for 1-2 hours at RT. Primary and secondary antibodies used in this study are summarized in **Table S5**. After one wash of 5 min with PBS, the slides were mounted with DAKO mounting medium (Agilent, S302380-2). Slides were left at RT to dry overnight and kept at 4ºC for long-term storage.

### Image Acquisition

Immunostaining images of organoids from D10 to D20 were acquired with an upright LSM 800 confocal microscope (Zeiss). Immunostaining images of D40 and D120 organoids were acquired with a Pannoramic FLASH 250 II digital Slide Scanner (3DHISTECH). Brightfield images of intact organoids in **Fig. 1B** and **Fig. S1** were acquired with a widefield microscope (AxioVert.A1, Zeiss GmbH) with a SONY Chameleon®3 CM3-U3-31S4M CMOS camera (Zeiss GmbH). Images of whole organoids in **Fig. 5A** and **Fig. S13A** were acquired with a Google Pixel 6. Immunostaining panels were prepared in Inkscape.

### Imaging data analysis

#### Quantification of organoid diameter at D10-20

Measurement of organoid diameter from brightfield images was performed in Fiji software using the line tool. When organoids presented non-circular (e.g. oval) morphology, the largest dimension was measured. Data matrices of quantifications were processed in R software v4.2.2 using dplyr (v1.0.9) and visualized using ggplot2 (v3.4.0). To aid visualization of growth dynamics, a trendline was added to the plot, using a smoothed conditional means function (ggplot2::geom_smooth, formula = ‘y ~ x’).

#### Quantification of organoid, rosette, and OTX2^+^ areas at D40

Demarcation of organoid, rosette, and OTX2^+^ areas was performed manually by drawing regions of interest (ROIs) in the CaseViewer software (3DHISTECH), using 2 or 3 slices per organoid. Data matrices of quantifications were processed in R software v4.2.2 using dplyr (v1.0.9) and visualized using ggplot2 (v3.4.0). Statistical analyses were performed in R software by ordinary one-way analysis of variance (ANOVA). The threshold for statistical significance was p < 0.05. Where indicated: 0 ≤ p < 0.001, ***; 0.001 ≤ p < 0.01, **; 0.01 ≤ p < 0.05, *; p ≥ 0.05, ns.

### Single-cell RNA sequencing

#### Generation of single-cell suspension of organoid cells

Organoids used for scRNAseq were harvested, cut into 2 or 3 pieces using two P10 pipette tips, and washed with DPBS-/-. Each individual organoid was incubated in 1.5 mL of Trypsin (Thermo Fisher Scientific, 15400054)/Accutase (Sigma-Aldrich, A6964) (1:1) containing 1 μL/mL of TURBO™ DNase (Thermo Fisher Scientific, AM2238, 2 U/μL) in a gentleMACS Dissociator (Miltenyi Biotec, 130-093-235) in the program NTDK1. Tubes with dissociated cells were briefly spun down and 1.5 mL of buffer (ice-cold DPBS-/- (Thermo Fisher Scientific) with 0.1% BSA (Sigma-Aldrich)) was added to the dissociated cells. The cell suspension was spun at 400 g for 5 min at 4 ºC. The supernatant was aspirated, leaving a margin of around 200 μL, and cells were resuspended in 700 μL of buffer. The suspension was filtered through a 70 μm strainer once, and through a FACS tube cap twice.

#### FACS sorting of viable cells

The viability dye DraQ7 (Biostatus, DR70250, 0.3 mM) was added at a concentration of 20 μL/mL and the suspension gently mixed with a P1000 pipette. 250k live cells of each individual organoid were FACS sorted using a 100 μm nozzle; singlets were gated based on forward and side scatter, and live cells based on negative excitation with an Alexa 700 filter.

#### Organoid multiplexing

The 10X Genomics 3’ CellPlex Kit was used to multiplex each individual organoid with a unique Cell Multiplexing Oligo (CMO), as described in the manufacturer’s protocol, except for the use of the aforementioned buffer formulation and 400 g in the spinning steps.

#### Library preparation

After multiplexing, live cells of each organoid were counted and pooled in equal numbers, normalized to the organoid with the lowest live cell count. The final pool was spun and resuspended in a small volume, and live cells were counted again. Two libraries were prepared, having been loaded with 40k and 50k live cells per channel (to give estimated recovery of 10k cells per channel) onto a Chromium Single Cell 3’ B Chip (10X Genomics, PN-1000073) and processed through the Chromium controller to generate single-cell GEMs (Gel Beads in Emulsion). ScRNAseq libraries were prepared with the Chromium Single Cell 3’ Library & Gel Bead Kit v.3 (10x Genomics, PN-1000075).

#### Library sequencing

The two libraries were pooled in a NovaSeq S4 flowcell (Illumina, together with other samples) and pair-end sequenced.

### Single-cell RNA data analysis

#### Data pre-processing

ScRNAseq reads were processed with Cell Ranger multi v6.0.1 (10X Genomics), using the prebuilt 10X GRCh38 reference refdata-gex-GRCh38-2020-A, and including introns. Further processing such as dimensionality reduction, clustering, and visualization of the scRNAseq data was performed in R software v4.2.2 with Seurat v4.2.0. No batch effect was detected between the two libraries, therefore they were processed jointly.

#### Identification of non-telencephalic clusters

Non-telencephalic cells were separated at clustering resolution 0.1 (“FindClusters”). Non-telencephalic clusters (clusters 3, 5 and 6) were removed for downstream analysis, based on absent/low FOXG1 expression.

#### Removal of low-quality cells

After exclusion of non-telencephalic cells, we used the Gruffi algorithm to identify and remove cells presenting a transcriptional signature of cellular stress (**Vértesy et al**., 2022). Cells with 800 to 6000 detected genes, and less than 6% mitochondrial and 30% ribosomal content were retained. Count data was log-normalized and scaled regressing out the number of genes and percentage of mitochondrial and ribossomal RNAs. Dimensionality reduction was performed using PCA on the top 2000 most variable genes, and the first 20 PCs were selected for the subsequent analysis.

#### Identification of telencephalic clusters

Clustering at resolution 1 and 0.5 yielded the most reliable separation by cell-types, based on the expression of known marker genes: dividing NPCs and RGs (clusters 3, 5, 7, 8, 12 at resolution 1), OPCs (cluster 10 at resolution 1), INs (cluster 9 at resolution 1), IPCs (cluster 4 at resolution 1), immature excitatory neurons (clusters 0, 1 and 2 at resolution 1; and clusters 0 and 6 at resolution 0.5), ExNs DL (cluster 1 at resolution 0.5) and ExNs UL (clusters 6 and 11 at resolution 1).

#### Data plotting

Two-dimensional representations were generated using uniform manifold approximation and projection (UMAP) (uwot v0.1.14). Data matrices of quantifications were processed in R software v4.2.2 using dplyr (v1.0.9) and visualized using ggplot2 (v3.4.0).

---

**Figure S1.**
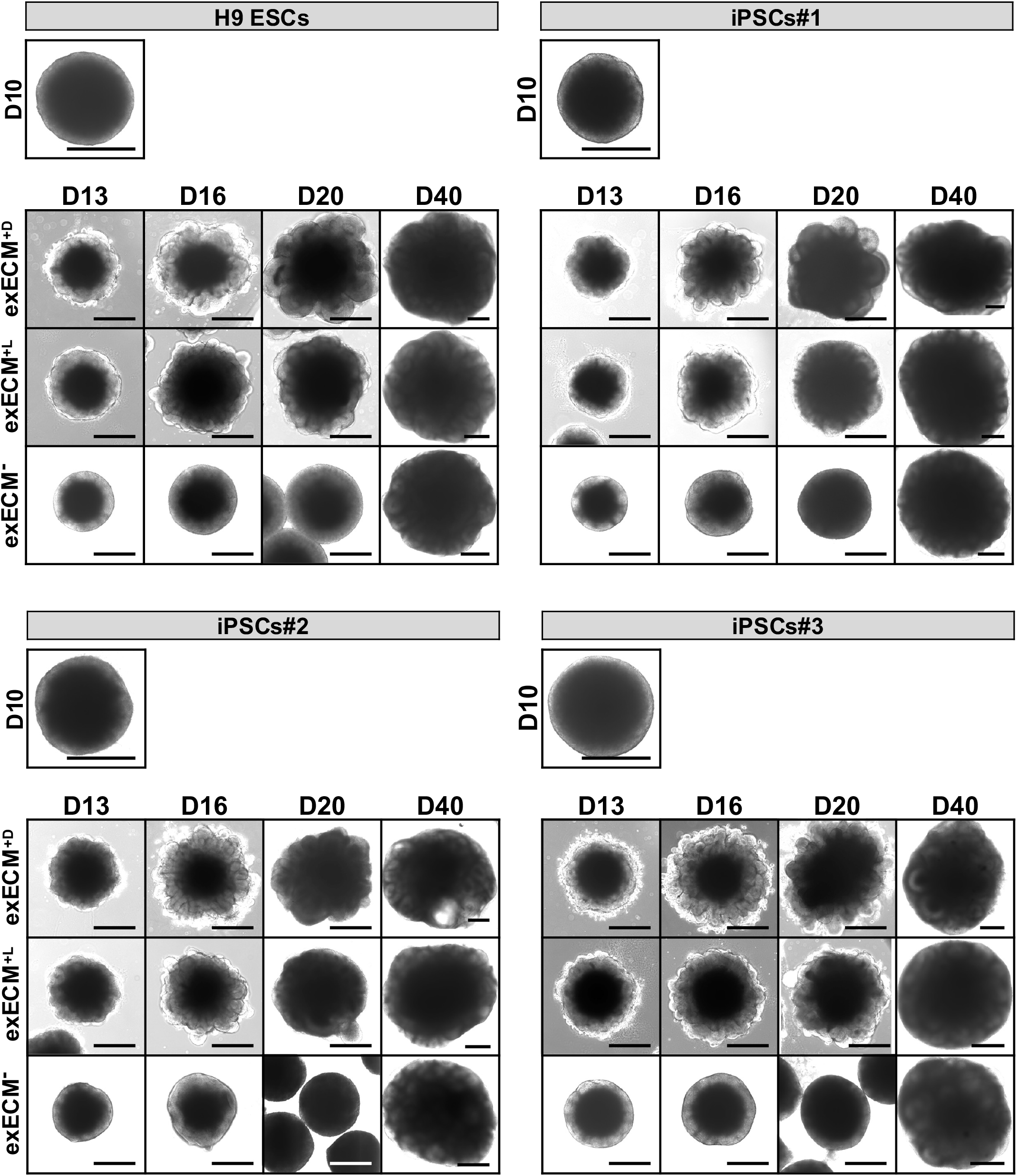
(Related to Fig. 1B) Organoid morphology at D10, D13, D16, D20, and D40 evolves in comparable ways in all cell lines. Morphological changes that are comparable across cell lines include outer brightening at D10, tissue budding in exECM^+D^ and exECM^+L^ conditions from D16, and formation of neural rosettes across conditions at D40. Scale bars: 500 μm.

**Figure S2.**
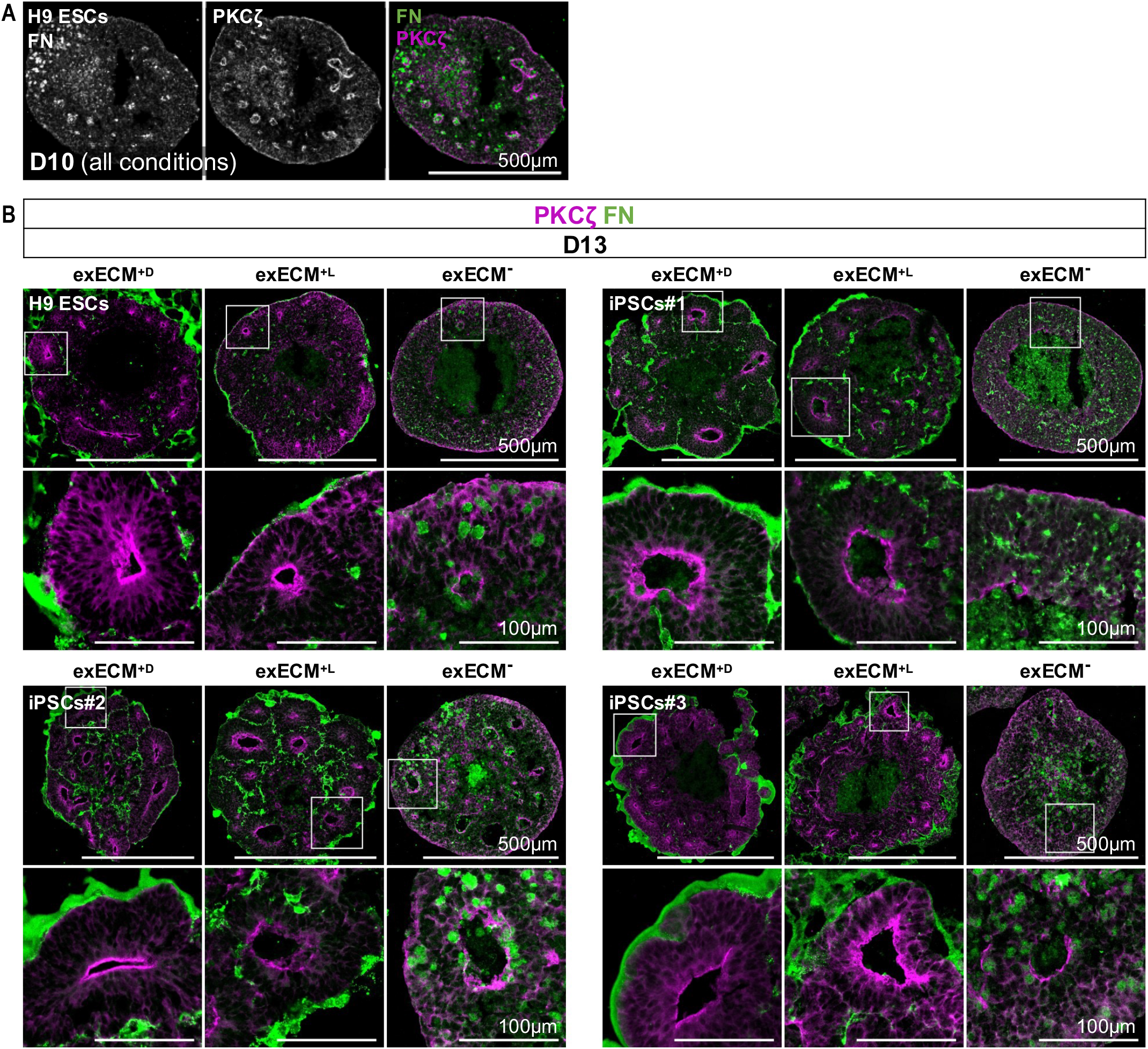
(Related to Fig. 3A-D) Comparison of the distribution of polarity proteins at D10 (A) and D13 (B) shows the quick action of exECM in tissue morphogenesis. Apical and basal domains are marked by PKCζ and FN, respectively. An apical-in/basal-out polarity axis is defined at D13 in exECM^+D^ and exECM^+L^ organoids, but still undefined in most exECM^-^ organoids. Bottom panels: magnification of inset.

**Figure S3.**
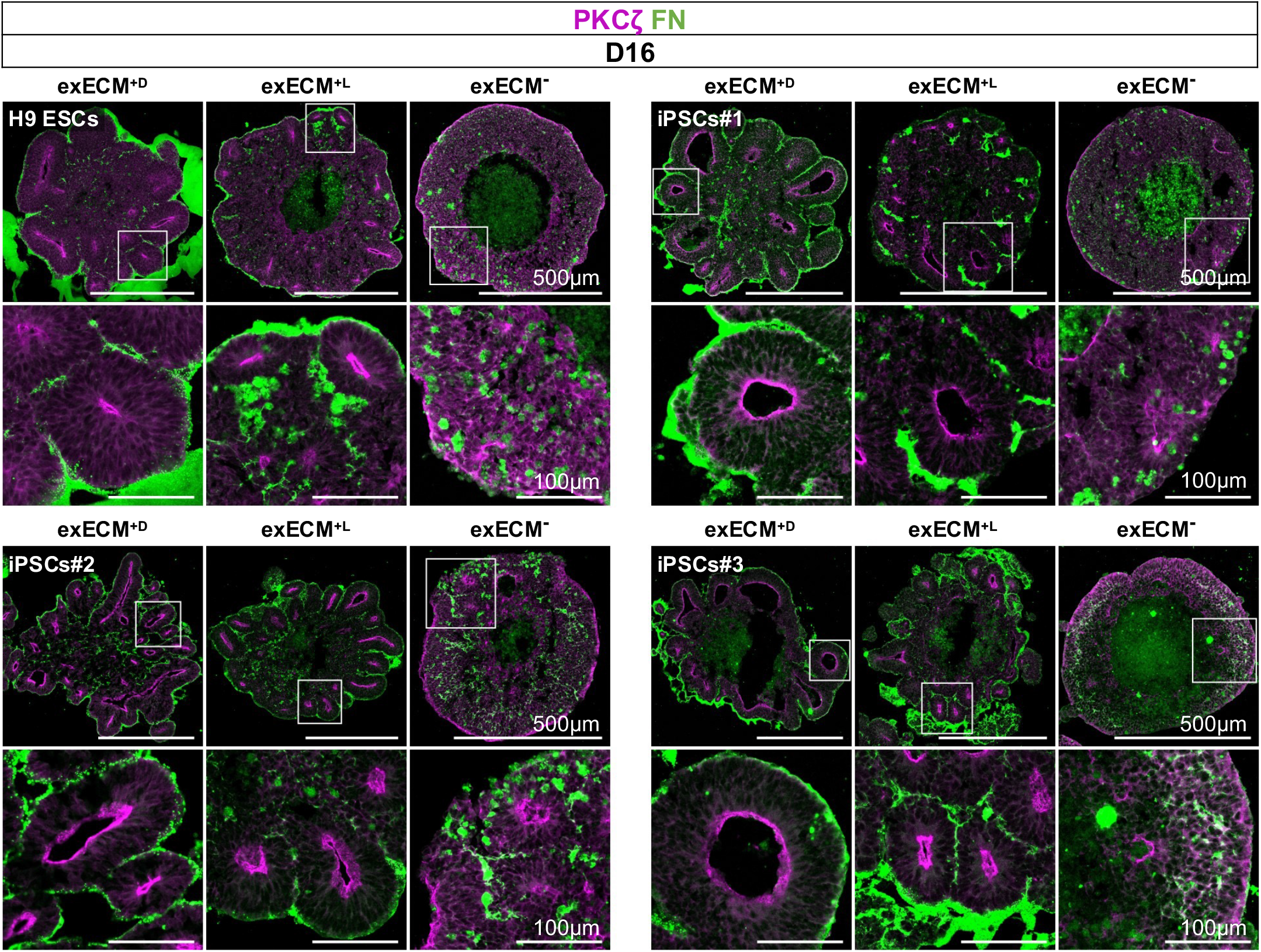
(Related to Fig. 3A-D) Apical-basal polarity axis at D16 differs between exECM^+^ and exECM^-^ conditions. Apical and basal domains are marked by PKCζ and FN, respectively. An apical-in/basal-out polarity axis is defined in exECM^+D^ and exECM^+L^ organoids, but still undefined in most exECM^-^ organoids. Bottom panels: magnification of inset.

**Figure S4.**
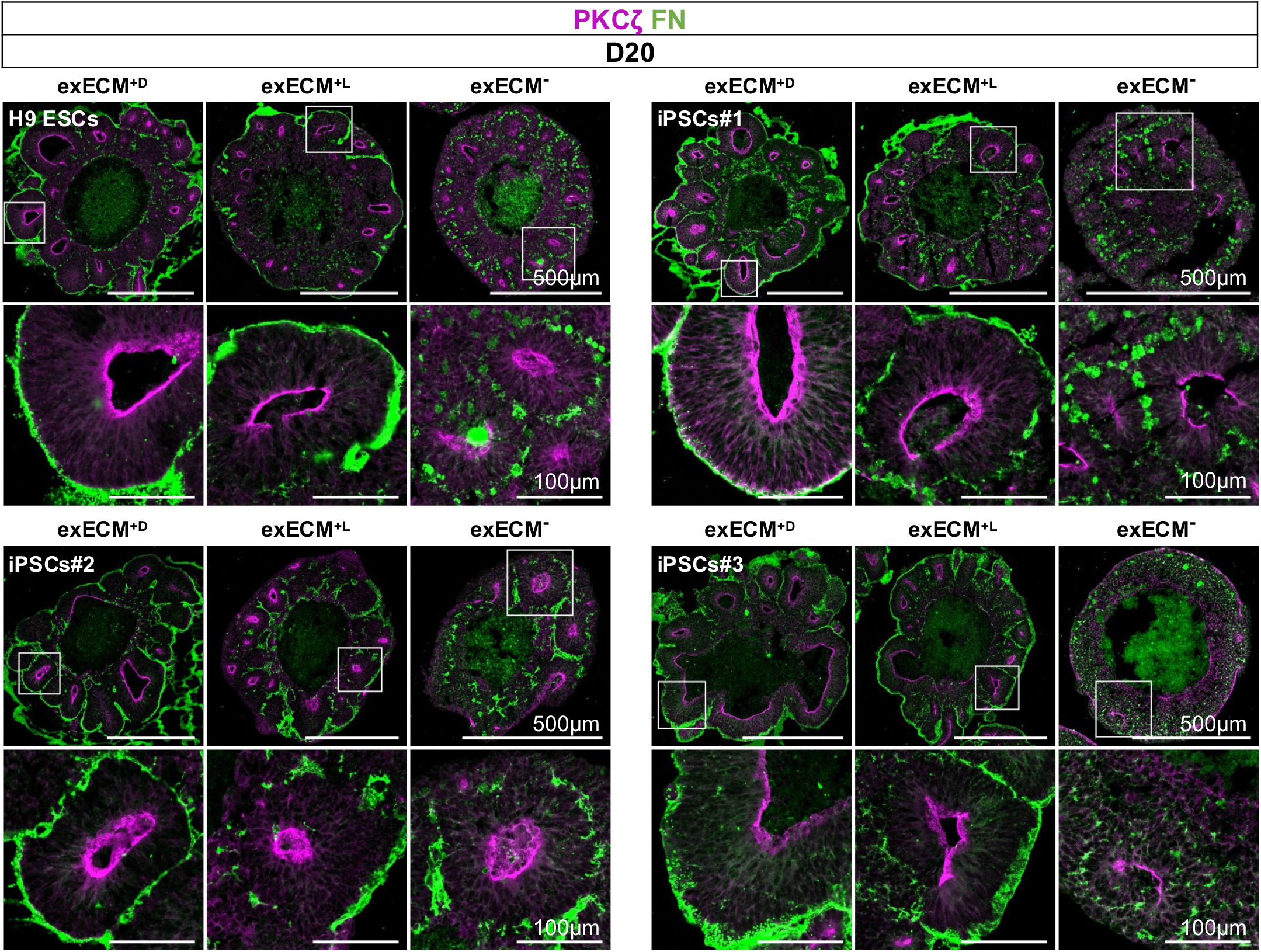
(Related to Fig. 3A-D) Apical-basal polarity axis at D20 is comparable between exECM^+^ and exECM^-^ conditions. Apical and basal domains are marked by PKCζ and FN, respectively. An apical-in/basal-out polarity axis is defined in all conditions. Bottom panels: magnification of inset.

**Figure S5.**
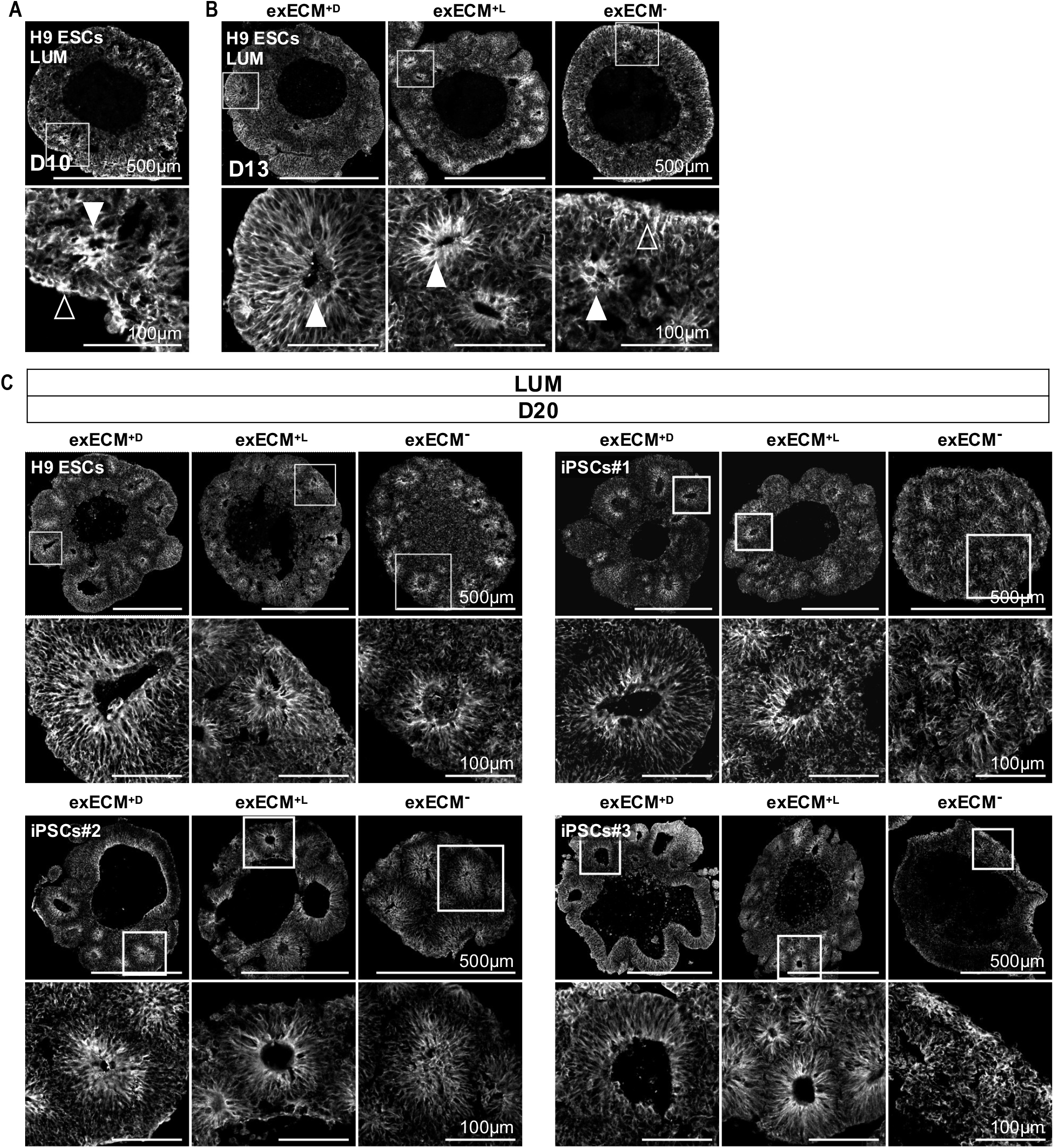
(Related to Fig. 3F) Lumican is endogenously produced by NPCs in all conditions. **(A)** At D10, LUM is abundantly expressed but presents disorganized tissue distribution. Apical localization to rosette ventricular zones is defined at D13 in exECM^+D^ and exECM^+L^ organoids **(B)** and at D20 in all conditions **(C)**. Arrowheads mark the location of LUM lining the organoid outer surface (Δ) or the ventricular zone of neural rosettes within the tissue (▲). Bottom panels: magnification of inset.

**Figure S6.**
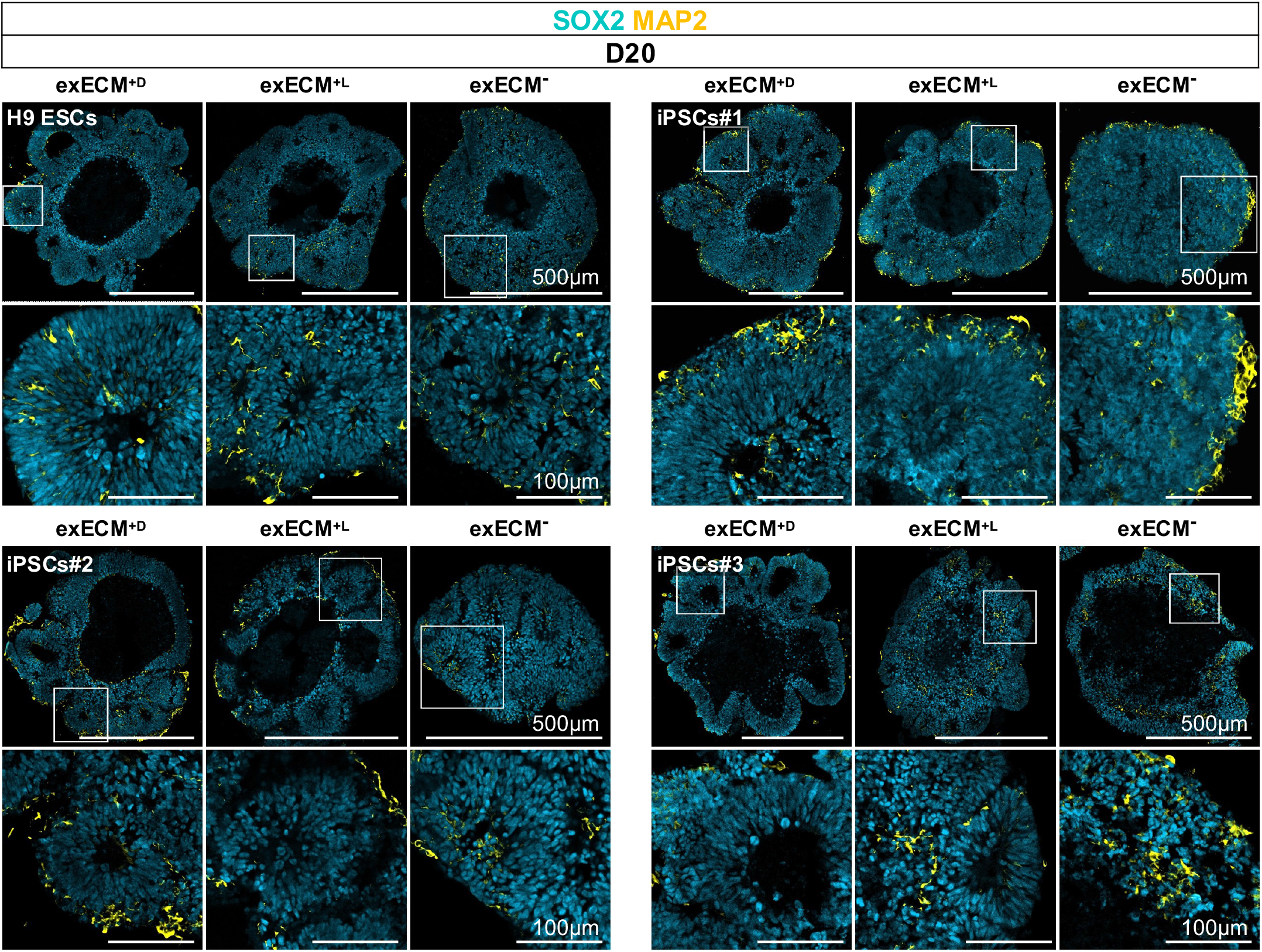
(Related to Fig. 4B) The onset of neurogenesis occurs at around D20. At D20, few scattered neurons (MAP2^+^) are seen mainly at the outer organoid surface or surrounding neural rosettes (SOX2^+^), in all conditions and cell lines. Bottom panels: magnification of inset.

**Figure S7.**
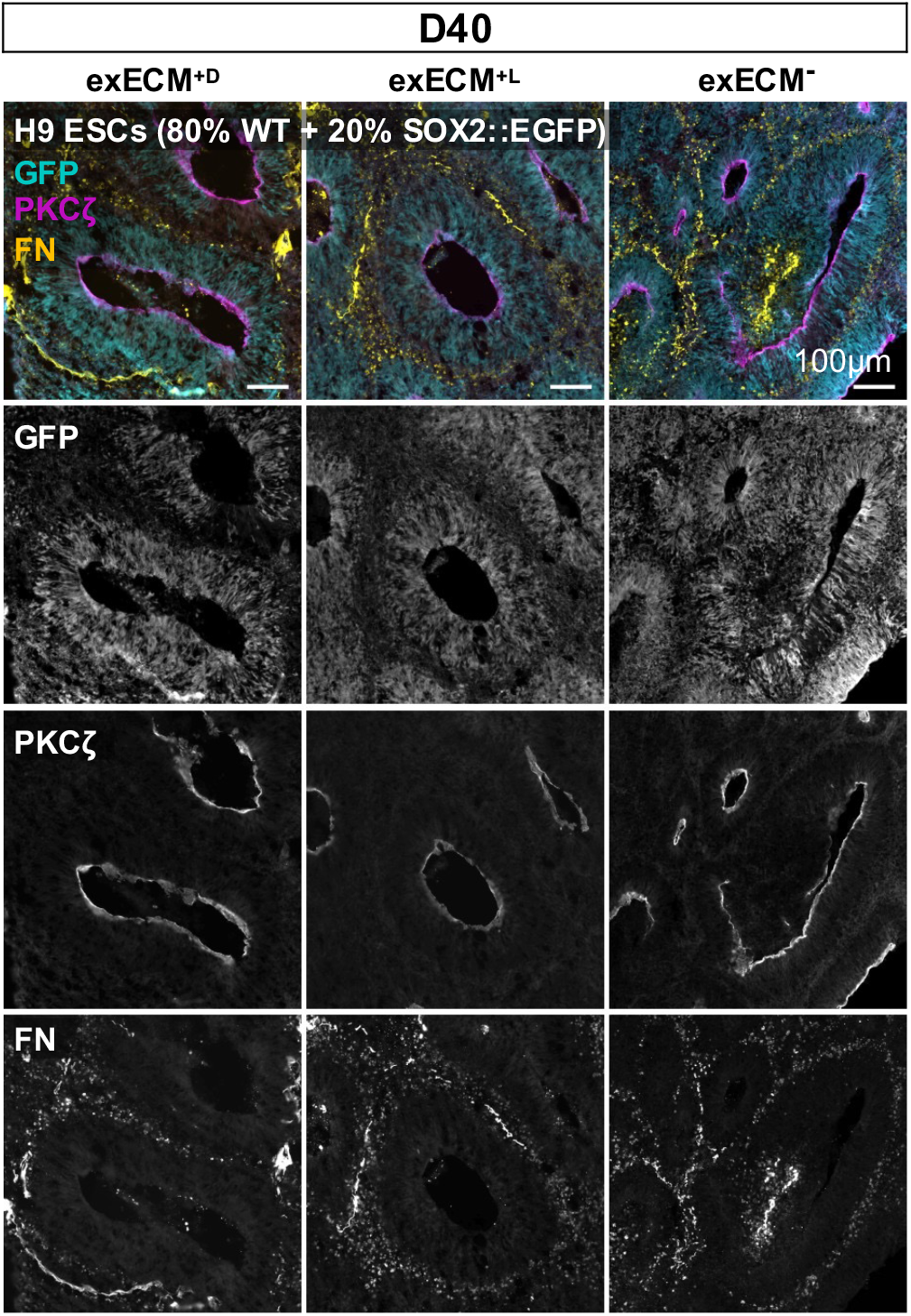
(Related to Fig. 4A-C) Rosette polarity and progenitor arrangement are comparable between conditions at D40. Neural rosettes show a PKCζ^+^ apical lumen and a surrounding FN^+^ basal region; SOX2^+^ neural progenitors are arranged radially around the rosette ventricular zone, as seen by GFP staining of organoids generated from 20% SOX2::EGFP and 80% WT H9 ESCs.

**Figure S8.**
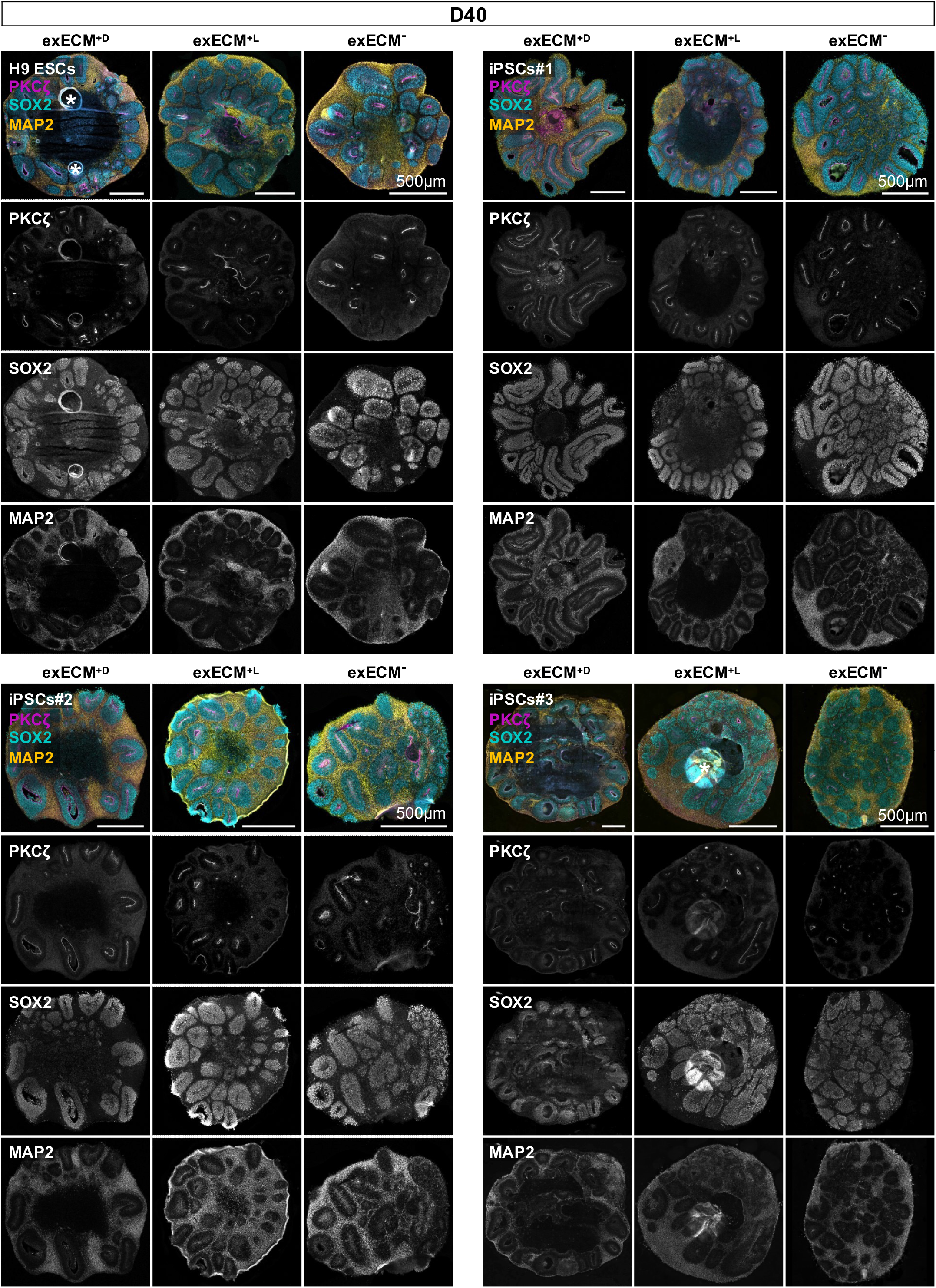
(Related to Fig. 4B) Organization of neural progenitors and neurons is comparable between conditions at D40. Organoids from all conditions and cell lines show abundant neural rosettes with PKCζ^+^ ventricular zone, and inside-out organization of SOX2^+^ neural progenitors and MAP2^+^ neurons. *: staining artifact.

**Figure S9.**
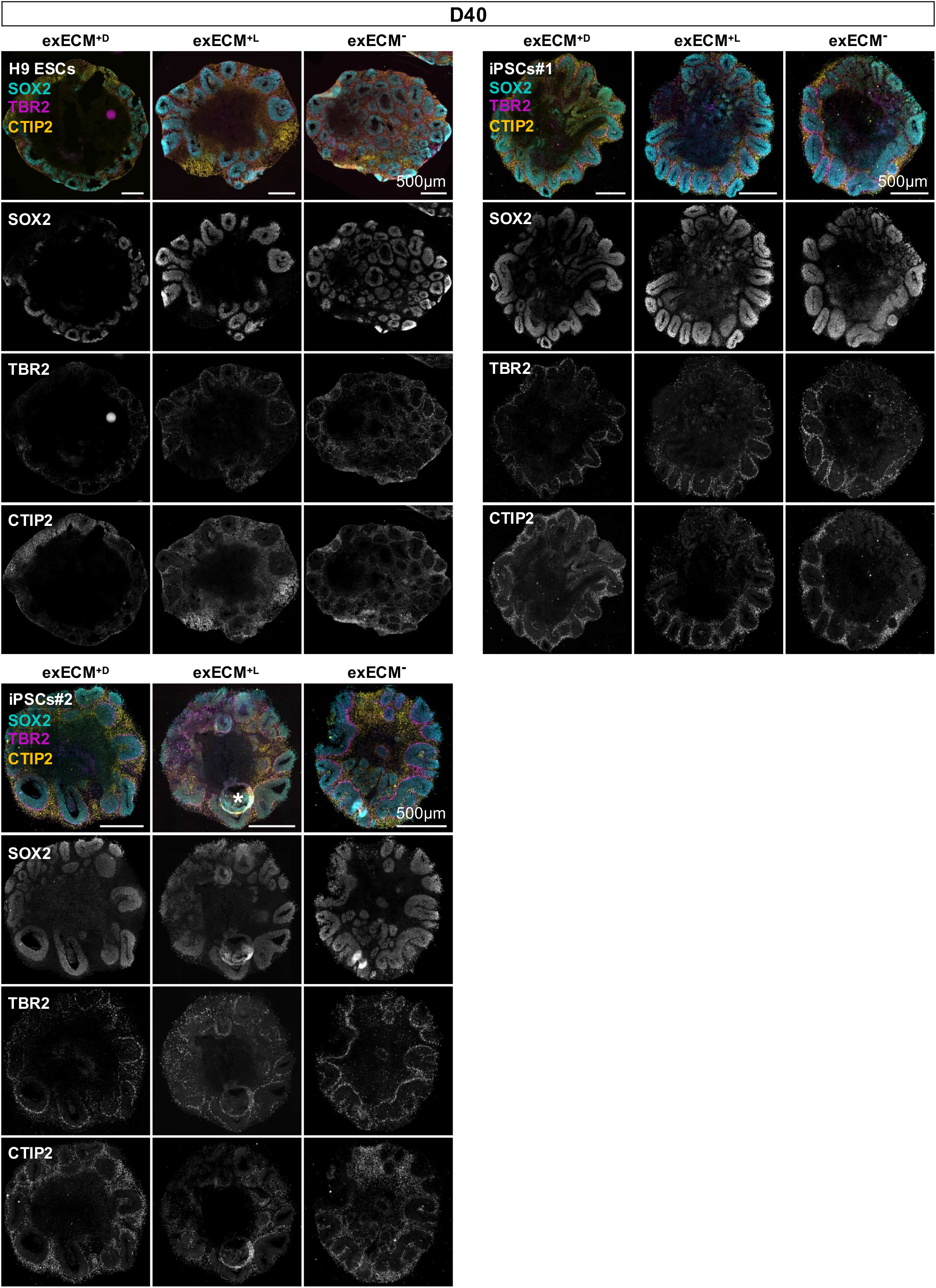
(Related to Fig. 4C) Tissue identity and organization of dorsal-cortical cell types is comparable between conditions at D40. Radial glia (SOX2^+^), dorsal intermediate progenitors (TBR2^+^) and early born excitatory neurons (CTIP2^+^) show layered arrangement in neural rosettes of organoids from all conditions and cell lines. *: staining artifact.

**Figure S10.**
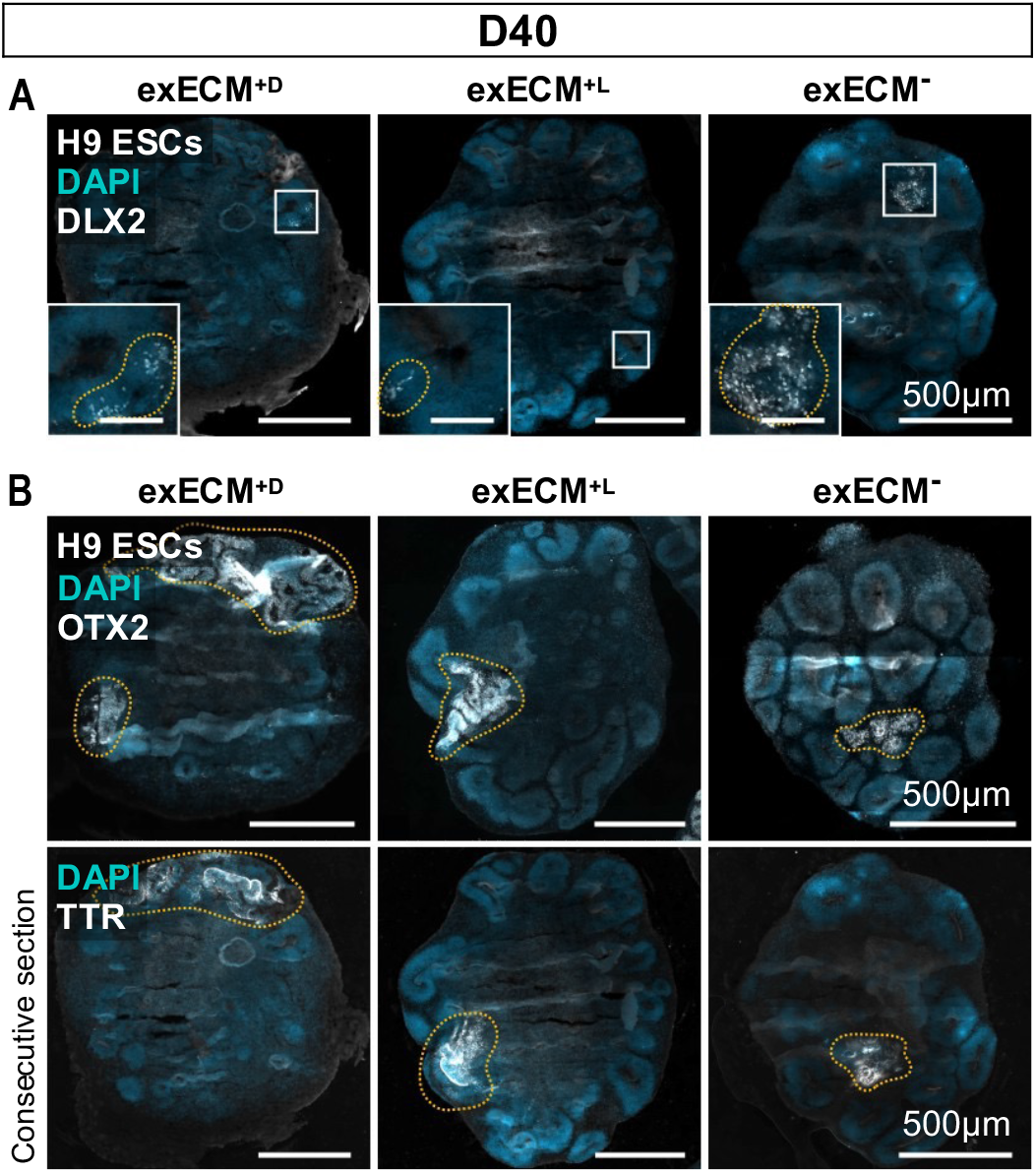
(Related to Fig. 4D and G) Small regions of mis-patterned cells are visible at D40. **(A)** At D40, small regions of DLX2^+^ ventral neural progenitor cells are seen in organoids from all conditions (demarcated by dashes). **(B)** Optic cup-like regions show convoluted morphology and are marked by the expression of OTX2 and TTR (shown in subsequent sections of the same organoid, demarcated by dashes).

**Figure S11.**
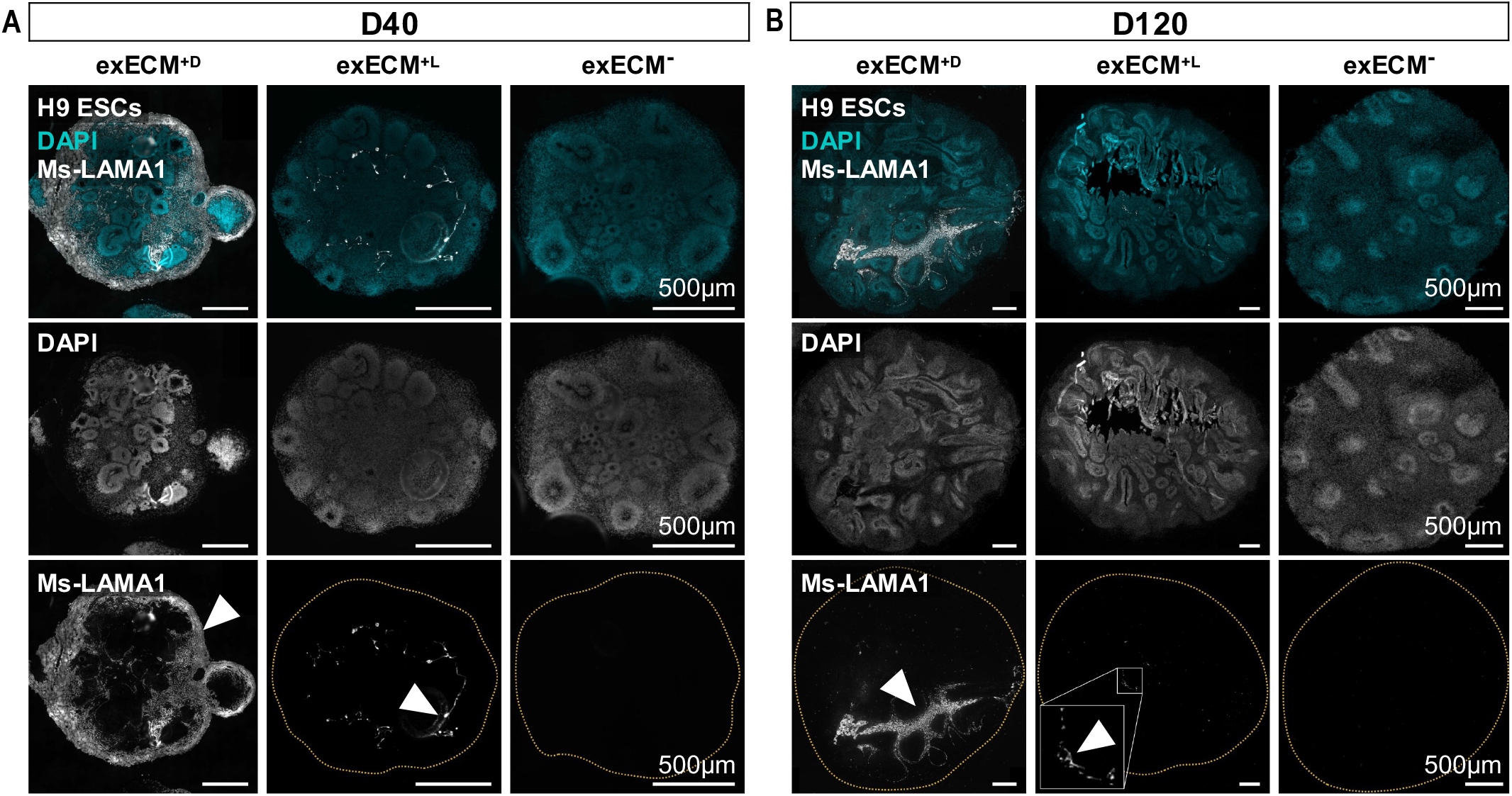
(Related to Figs. 3E, 4 and 5) Matrigel shows long-term permanence within the organoid tissue. **(A)** At D40, exECM^+D^ organoids remain encapsulated by Matrigel (marked by Ms-LAMA1, ▲); and Matrigel remnants are visible within the tissue of exECM^+L^ organoids. **(B)** At D120, large Ms-LAMA1^+^ regions are visible in exECM^+D^ organoids, when Matrigel has been engulfed by the growing tissue; Matrigel remnants are still present in exECM^+L^organoids. ExECM^-^ organoids show complete absence of Ms-LAMA1 staining, as expected.

**Figure S12.**
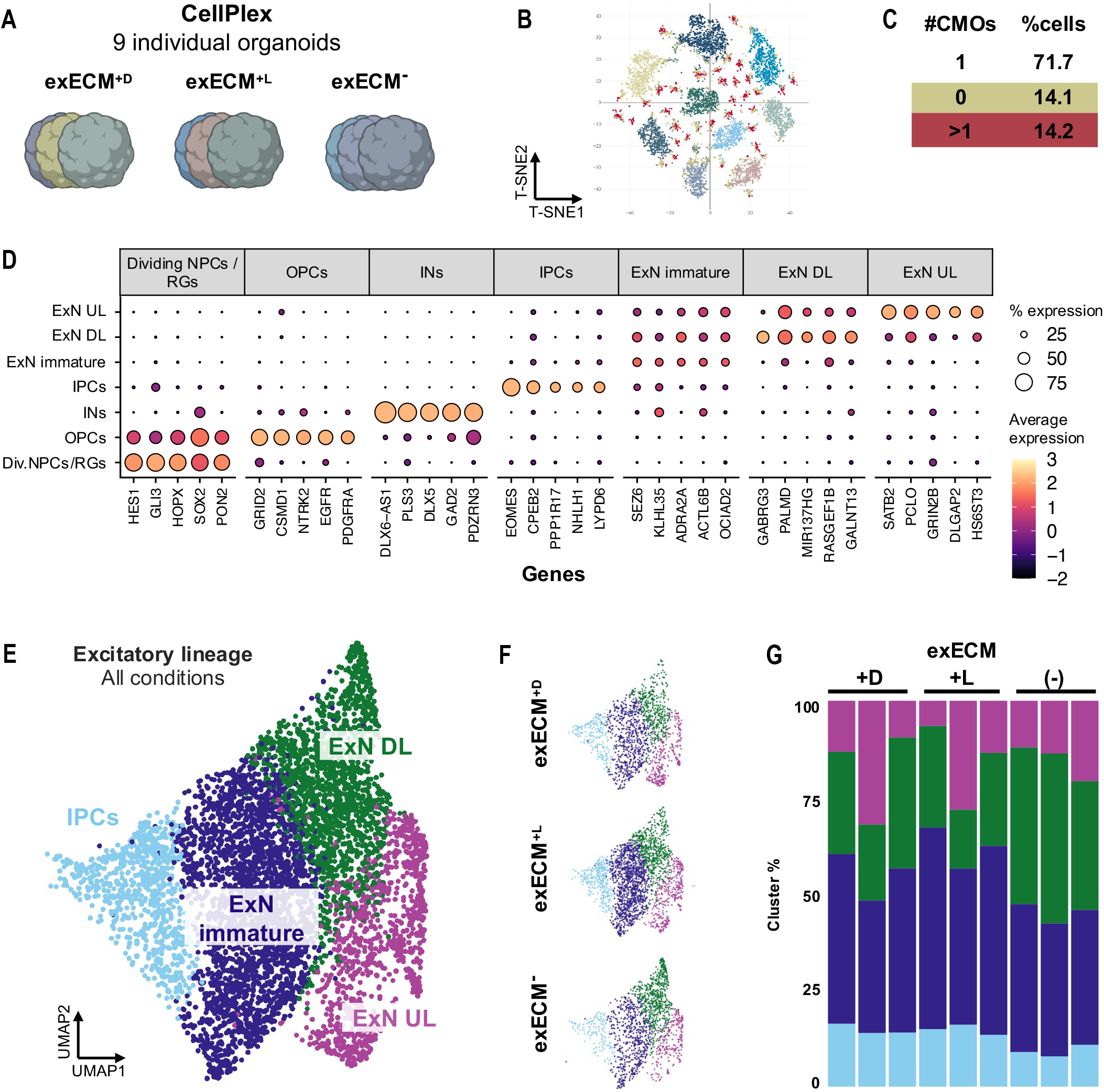
(Related to Fig. 5A-H) Details of scRNAseq analysis. **(A)** Nine individual organoids (3 per exECM condition) were multiplexed with unique molecular identifiers (Cell Multiplexing Oligos, CMOs). **(B)** T-SNE projection of cells retrieved after sequencing, based on recovered CMOs. **(C)** Demultiplexing of used CMOs allows the assignment of individual cells to the respective organoid of origin. More than 70 % of cells (corresponding to 6714 cells) were assigned to an individual organoid; cells tagged with zero (12.9 %) or more than one CMO (15%) were excluded from the downstream analysis. Here, one of two sequenced libraries is shown; in the second library, about 56 % of cells (7756 cells) were assigned to a unique CMO. **(D)** Top 5 cluster markers of telencephalic clusters: dividing NPCs/radial glia progenitors (RGs), oligodendrocyte precursor cells (OPCs), interneurons (INs), intermediate progenitor cells (IPCs), immature excitatory neurons (ExN), deep-layer excitatory neurons (ExNs DL) and upper-layer ExNs (ExNs UL). **(E)** Subset of excitatory neuron lineage in UMAP projection after filtering and exclusion of non-telencephalic clusters; selected clusters: IPCs, immature exNs, deep-layer ExNs and upper-layer ExNs. **(F)** UMAP projection of the same dataset for each individual exECM condition. **(G)** Calculation of the percentage of each cluster per condition and per organoid.

**Figure S13.**
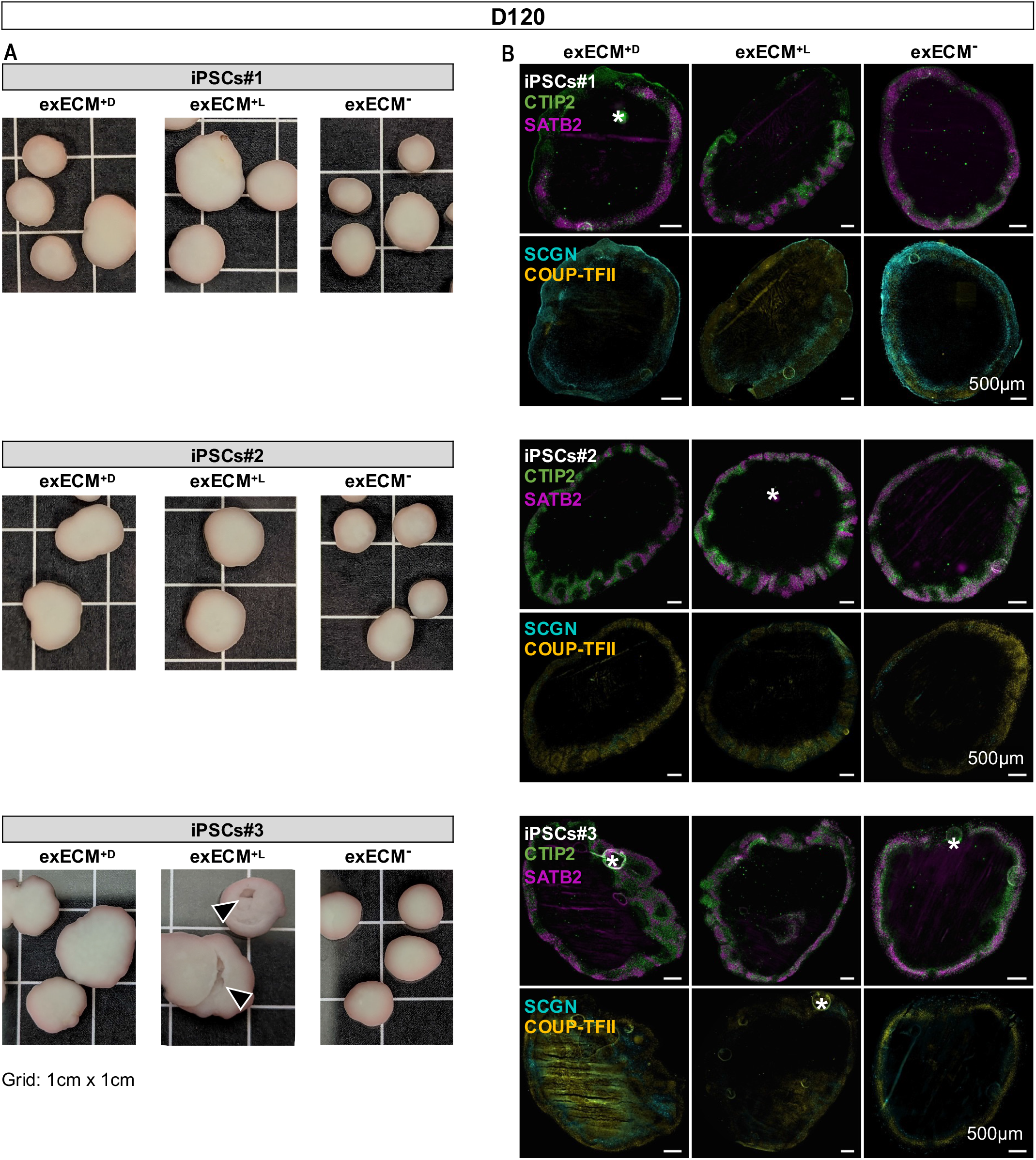
(Related to Fig. 5I) Organoids cultured for 120 days show size dependent on exECM conditions, but comparable cell types and tissue morphology. **(A)** Although organoid size is variable at D120, exECM^-^ organoids are generally smaller than exECM^+D^ or exECM^+L^ organoids. Overgrowth may cause tissue damage, as seen in iPSCs#3-derived exECM^+L^ organoids (▲). **(B)** Tissue immunostaining of organoids at D120 shows abundant deep- and upper-layer neurons (CTIP2^+^ and SATB2^+^, respectively), as well as less abundant populations of interneurons (SCGN^+^ and COUP-TFII^+^) across all conditions and cell lines. *: staining artifacts.

## Supplementary Tables

**Table S1.** (Related to Fig. 4D) Statistical analysis on the organoid diameter at D40. Ordinary one-way analysis of variance (ANOVA); 0 ≤ p < 0.001, ***; 0.001 ≤ p < 0.01, **; 0.01 ≤ p < 0.05, *; p ≥ 0.05, ns. Drop, exECM^+D^; Liq, exECM^+L^; Null, exECM^-^.

| Cell Line | Condition | Mean difference | Lower range | Upper range | Adj. p-val. | Significance |
| --- | --- | --- | --- | --- | --- | --- |
| H9 ESCs | Liq-Drop | -784.444 | -1122.24 | -446.65 | 3.28E-06 | *** |
| H9 ESCs | Null-Drop | -1156.16 | -1493.96 | -818.369 | 3.85E-10 | *** |
| H9 ESCs | Null-Liq | -371.719 | -719.306 | -24.1318 | 0.033663 | * |
| iPSCs#1 | Liq-Drop | -164.933 | -604.69 | 274.8244 | 0.63151 | ns |
| iPSCs#1 | Null-Drop | -573.515 | -994.551 | -152.48 | 0.005734 | * |
| iPSCs#1 | Null-Liq | -408.583 | -848.34 | 31.17478 | 0.072836 | ns |
| iPSCs#2 | Liq-Drop | -569.957 | -805.451 | -334.463 | 1.46E-06 | *** |
| iPSCs#2 | Null-Drop | -831.263 | -1066.76 | -595.769 | 1.63E-10 | *** |
| iPSCs#2 | Null-Liq | -261.306 | -493.204 | -29.4079 | 0.02394 | * |
| iPSCs#3 | Liq-Drop | -875.703 | -1057.04 | -694.368 | 1.81E-14 | *** |
| iPSCs#3 | Null-Drop | -1015.06 | -1185.78 | -844.346 | 8.77E-15 | *** |
| iPSCs#3 | Null-Liq | -139.36 | -307.304 | 28.58335 | 0.124131 | ns |

**Table S2.** (Related to Fig. 4E) Statistical analysis on the median rosette area in D40 organoids. Ordinary one-way analysis of variance (ANOVA); 0 ≤ p < 0.001, ***; 0.001 ≤ p < 0.01, **; 0.01 ≤ p < 0.05, *; p ≥ 0.05, ns. Drop, exECM^+D^; Liq, exECM^+L^; Null, exECM^-^.

| Cell Line | Condition | Mean difference | Lower range | Upper range | Adj. p-val. | Significance |
| --- | --- | --- | --- | --- | --- | --- |
| H9 ESCs | Liq-Drop | -0.0113 | -0.02098 | -0.00162 | 0.019008 | * |
| H9 ESCs | Null-Drop | -0.01344 | -0.02267 | -0.0042 | 0.003009 | * |
| H9 ESCs | Null-Liq | -0.00213 | -0.01116 | 0.006894 | 0.832851 | ns |
| iPSCs#1 | Liq-Drop | -0.01607 | -0.02613 | -0.00601 | 0.001512 | * |
| iPSCs#1 | Null-Drop | -0.02433 | -0.03381 | -0.01485 | 3.68E-06 | *** |
| iPSCs#1 | Null-Liq | -0.00826 | -0.01832 | 0.001795 | 0.121686 | ns |
| iPSCs#2 | Liq-Drop | -0.00187 | -0.00909 | 0.005349 | 0.805481 | ns |
| iPSCs#2 | Null-Drop | -0.01119 | -0.01841 | -0.00397 | 0.001425 | * |
| iPSCs#2 | Null-Liq | -0.00932 | -0.01593 | -0.00271 | 0.003839 | * |
| iPSCs#3 | Liq-Drop | -0.02376 | -0.02985 | -0.01767 | 0 | *** |
| iPSCs#3 | Null-Drop | -0.02784 | -0.03405 | -0.02164 | 0 | *** |
| iPSCs#3 | Null-Liq | -0.00408 | -0.01003 | 0.001871 | 0.23475 | ns |

**Table S3.** (Related to Fig. 4F) Statistical analysis on the area of OTX2+ tissue in D40 organoids. Ordinary one-way analysis of variance (ANOVA); 0 ≤ p < 0.001, ***; 0.001 ≤ p < 0.01, **; 0.01 ≤ p < 0.05, *; p ≥ 0.05, ns. Drop, exECM^+D^; Liq, exECM^+L^; Null, exECM^-^.

| Cell Line | Condition | Mean difference | Lower range | Upper range | Adj. p-val. | Significance |
| --- | --- | --- | --- | --- | --- | --- |
| H9 ESCs | Liq-Drop | -11.7291 | -19.6897 | -3.76851 | 0.002662 | * |
| H9 ESCs | Null-Drop | -13.5799 | -21.1738 | -5.98592 | 0.000291 | *** |
| H9 ESCs | Null-Liq | -1.85073 | -9.27589 | 5.574432 | 0.816015 | ns |
| iPSCs#1 | Liq-Drop | -7.40224 | -14.297 | -0.50753 | 0.033693 | * |
| iPSCs#1 | Null-Drop | -8.83626 | -15.3367 | -2.33586 | 0.006508 | * |
| iPSCs#1 | Null-Liq | -1.43402 | -8.32873 | 5.460694 | 0.862659 | ns |
| iPSCs#2 | Liq-Drop | -21.2544 | -31.0578 | -11.451 | 1.21E-05 | *** |
| iPSCs#2 | Null-Drop | -29.4243 | -39.2277 | -19.6209 | 1.33E-08 | *** |
| iPSCs#2 | Null-Liq | -8.16991 | -17.148 | 0.808163 | 0.081067 | ns |
| iPSCs#3 | Liq-Drop | -0.45795 | -10.5524 | 9.636541 | 0.993507 | ns |
| iPSCs#3 | Null-Drop | -14.1598 | -24.4498 | -3.86975 | 0.004382 | * |
| iPSCs#3 | Null-Liq | -13.7018 | -23.6841 | -3.71954 | 0.004492 | * |

**Table S4.**
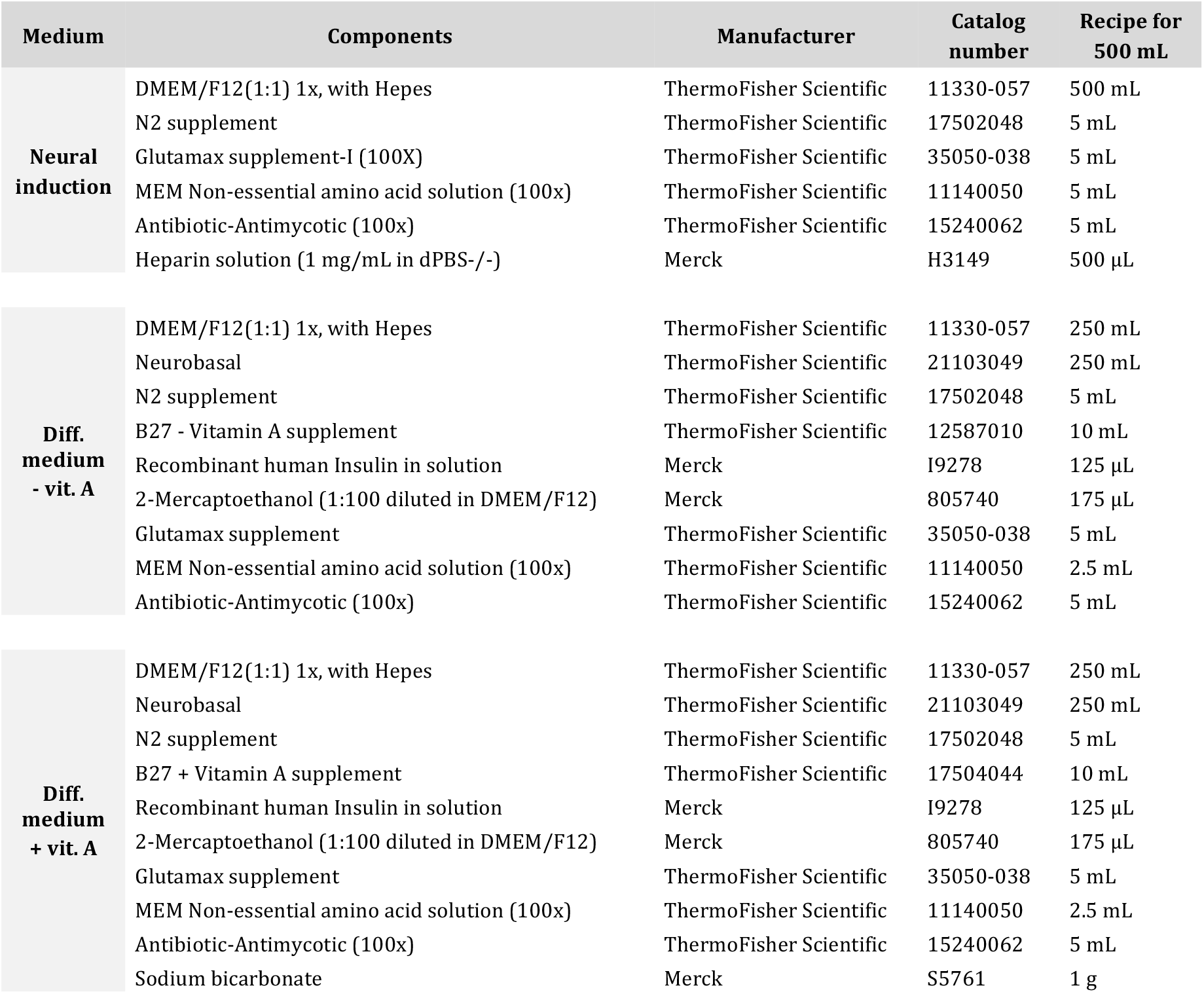
Medium composition for telencephalic organoid culture. Medium components and recipe for 500 mL of Neural Induction medium (NI), Differentiation medium without vitamin A (Diff-A), and Differentiation medium with Vitamin A (Diff+A).

**Table S5.** Antibodies used in this study.

| Primary antibodies |  |  |  |
| --- | --- | --- | --- |
| Species | Antigen | Manufacturer | Catalog# |
| Chicken | GFP | Aves Labs | GFP-1020 |
| Mouse | PKCζ | Santa Cruz Biotechnology | sc-17781 |
| Rabbit | PKCζ | Santa Cruz Biotechnology | sc-216 |
| Mouse | Fibronectin | R&D Systems | MAB1918 |
| Rat | LAMA1 | R&D Systems | MAB4656 |
| Rabbit | Lumican | Abcam | ab168348 |
| Mouse | SOX2 | R&D Systems | MAB2018 |
| Goat | SOX2 | R&D Systems | AF2018 |
| Chicken | MAP2 | Thermo Fisher Scientific | PA1-10005 |
| Rabbit | TBR2 | Abcam | ab23345 |
| Sheep | TBR2 | R&D Systems | AF6166 |
| Rat | CTIP2 | Abcam | ab18465 |
| Mouse | SATB2 | Abcam | ab51502 |
| Goat | OTX2 | Neuromics | GT15095 |
| Sheep | TTR | AbD Serotec | ahp 1837 |
| Mouse | DLX2 | Santa Cruz Biotechnology | sc-393879 |
| Rabbit | SCGN | Sigma-Aldrich | HPA006641 |
| Mouse | COUP-TFII | R&D Systems | pp-h7147-00 |

| Secondary antibodies |  |  |  |
| --- | --- | --- | --- |
| Species | Antigen | Manufacturer | Catalog# |
| Alexa Fluor 488 Donkey anti-rabbit |  | Invitrogen | A21206 |
| Alexa Fluor 488 Donkey anti-mouse |  | Invitrogen | A21202 |
| Alexa Fluor 488 Donkey anti-chicken |  | Jackson ImmunoResearch | 703-545-155 |
| Alexa Fluor 488 Donkey anti-goat |  | Invitrogen | A11055 |
| Alexa Fluor 568 Donkey anti-rabbit |  | Invitrogen | A10042 |
| Alexa Fluor 568 Donkey anti-mouse |  | Invitrogen | A10037 |
| Alexa Fluor 568 Donkey anti-goat |  | Invitrogen | A11057 |
| Alexa Fluor 647 Donkey anti-rabbit |  | Invitrogen | A31573 |
| Alexa Fluor 647 Donkey anti-mouse |  | Invitrogen | A31571 |
| Alexa Fluor 647 Donkey anti-goat |  | Invitrogen | A21447 |
| Alexa Fluor 647 Donkey anti-rat |  | Jackson ImmunoResearch | 712-605-150 |

